# Efficient Retroelement-Mediated DNA Writing in Bacteria

**DOI:** 10.1101/2020.02.21.958983

**Authors:** Fahim Farzadfard, Nava Gharaei, Robert J. Citorik, Timothy K. Lu

## Abstract

The ability to efficiently and dynamically change information stored in genomes would enable powerful strategies for studying cell biology and controlling cellular phenotypes. Current recombineering-mediated DNA writing platforms in bacteria are limited to specific laboratory conditions, often suffer from suboptimal editing efficiencies, and are not suitable for *in situ* applications. To overcome these limitations, we engineered a retroelement-mediated DNA writing system that enables efficient and precise editing of bacterial genomes without the requirement for target-specific elements or selection. We demonstrate that this DNA writing platform enables a broad range of applications, including efficient, scarless, and *cis*-element-independent editing of targeted microbial genomes within complex communities, the high-throughput mapping of spatial information and cellular interactions into DNA memory, and the continuous evolution of cellular traits.

**One Sentence Summary:** Highly-efficient, dynamic, and conditional genome writers are engineered for DNA memory, genome engineering, editing microbial communities, high-resolution mapping of cellular connectomes, and modulating cellular evolution.

## Main Text

Genomic DNA is an evolvable functional memory that records the history of adaptive changes over evolutionary timescales. DNA writing platforms that enable efficient and targeted modifications of genomic DNA are essential for studying and engineering living cells, with many applications ranging from the recording of cellular lineages and transient molecular events into permanent DNA records to cellular computation (Farzadfard and Lu, 2018). An ideal precise DNA writer (a genetically-encoded device for the targeted editing of DNA in living cells) would enable one to introduce any desired mutation to any desired genomic target with high efficiency and without the requirement for specific *cis*-encoded elements or the generation of double-strand DNA breaks. Despite many advances in recent years in DNA writing technologies, existing platforms in bacteria (Costantino and Court, 2003; Datsenko and Wanner, 2000; Farzadfard et al., 2019; Pines et al., 2015; Swingle et al., 2010; Wang et al., 2009; Yu et al., 2000) are not ideal for certain applications (Table S1). For example, recombineering-based approaches enable targeted, small modification of bacterial genomes but 1) they are restricted to specific conditions in which efficient transformation is possible, 2) are often limited by suboptimal editing rates, and 3) are not applicable to complex environments, such as bacterial communities (Costantino and Court, 2003; Wang et al., 2009; Yu et al., 2000). In addition, recombineering events cannot be linked to cellular regulatory networks and thus cannot be used for continuous and dynamic manipulation of cellular phenotypes, autonomous recording of cellular events histories, or evolutionary genome engineering. Although recombineering efficiencies have been improved by using CRISPR-Cas9 counter-selection (Jiang et al., 2013; Ronda et al., 2016), this strategy requires the presence of *cis*-encoded elements (i.e., the PAM domain) on the target and could also induce cytotoxic double-stranded breaks (Citorik et al., 2014; Cui and Bikard, 2016), which may limit the application space. Newer CRISPR-based DNA writing technologies such as base editing (Gaudelli et al., 2017b; Komor et al., 2016) and prime editing (Anzalone et al., 2019) have addressed some of these limitations. However, these technologies are currently limited as base editing can generate only a limited spectrum of mutations and the applicability of prime editing in bacteria is yet to be demonstrated.

To circumvent some of the above-mentioned limitations, we previously developed SCRIBE (Synthetic Cellular Recorders Integrating Biological Events), a retroelement-mediated precise DNA writing platform for conditional and targeted editing of bacterial genomes (Farzadfard and Lu, 2014). In this system, single-stranded DNAs are expressed intracellularly from an engineered retroelement (retron) cassette via reverse transcription and recombined into homologous sites on the genome by Beta-mediated recombination. The moderate recombination rate (∼10^-4^ recombination events per generation) achieved by the original SCRIBE system enables recording of the duration and magnitude of exposure of input(s) in the form of mutations that accumulate in the genomic DNA of bacterial populations, thus facilitating conversion of transcriptional signals into DNA memory. However, this level of recombination is not adequate for many applications that require a much more efficient DNA writing system.

In the present study, we sought to identify cellular factors that limit the recombination efficiency of retroelement-mediated recombination in *Escherichia coli* (*E. coli*). By systematically investigating these factors, we significantly improved SCRIBE efficiency and created HiSCRIBE (High-efficiency SCRIBE), a genetically-encoded precise DNA writing system that enables autonomous, dynamic, and transcriptionally controlled modification of bacterial genomes with high efficiency. HiSCRIBE writers achieve up to ∼100% editing efficiency in a scarless fashion, without generation of double-strand DNA breaks, or the requirement for the presence of cis-encoded sequences on the target, or selection.

We demonstrated the utility of this DNA writing platform for multiple applications (Fig. 1). Specifically, we showed that HiSCRIBE can be introduced into cells via different delivery mechanisms, including transduction and conjugation, enabling efficient and specific genome writing in bacteria within communities, which is not feasible with traditional oligo-mediated recombineering approaches. Furthermore, we demonstrated that efficient and precise DNA writing can be used to record transient spatial information (such as cell-cell interactions that happen during conjugation events in bacterial populations) into genomic DNA, allowing one to reduce multidimensional interactomes into a one-dimensional DNA sequence space, thus facilitating the study of complex cellular interactions within cell communities. Finally, we showed that when combined with a continuous delivery system and appropriate selection or screening, HiSCRIBE DNA writers can be used for the continuous optimization of a trait of interest. We envision that this highly-efficient DNA writing technology unlocks new avenues for the study of bacterial physiology and the dynamic engineering of cellular phenotypes.

**Figure 1.**
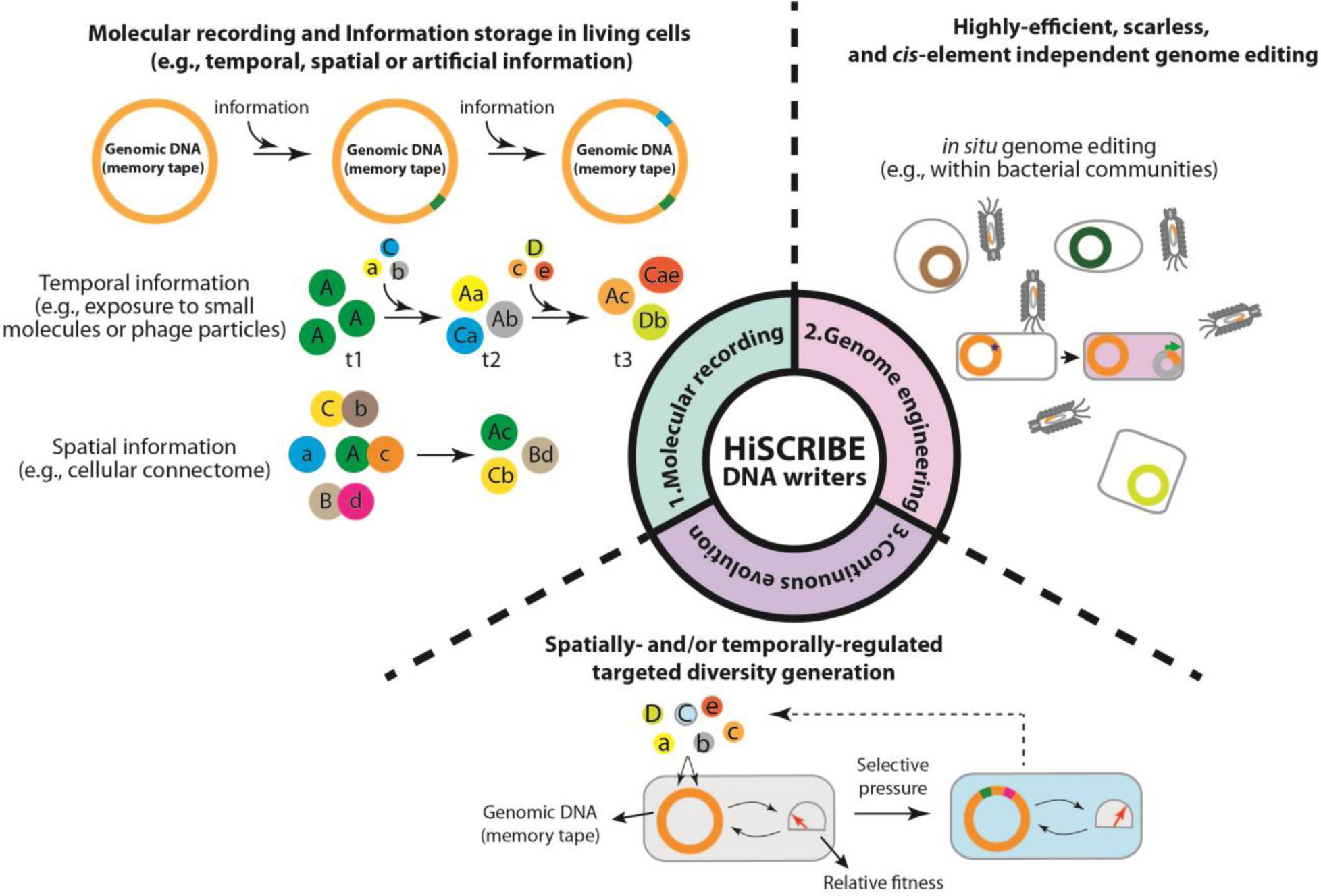
Distinctive applications enabled by high-efficiency SCRIBE (HiSCRIBE) DNA writers. (1) Recording spatiotemporal information into genomic DNA by writing unique barcodes into genomic DNA. Temporal information is recorded by introducing unique barcodes (letters in small circles) into the genomic DNA of individual cells (big circles) in response to incoming signals, either by conditional writing using an inducible promoter or by direct transfer of DNA from a mobilizable DNA element followed by writing of the barcode in the genome. Spatial information is recorded by a barcode joining strategy, where barcodes from interacting partners are brought together upon the interaction between the partners. (2) Highly efficient and scarless genome editing without the requirement for double-strand DNA breaks and target-specific *cis*-encoded elements enables genome editing within bacterial communities. (3) Spatially or temporally regulated diversity generation can be coupled to continuous selection for the continuous evolution of traits of interest.

## Optimizing SCRIBE for Molecular Recording and Highly-efficient Genome Writing

The SCRIBE platform opens up the entire genomic space for dynamic and precise DNA writing, as it does not require the presence of *cis*-encoded elements on a target. However, the moderate writing efficiency of the original system (10^-4^ events/generation) (Farzadfard and Lu, 2014) limits its utility to population-level molecular recording and makes it unsuitable for many applications that could benefit from higher DNA writing efficiencies (Farzadfard and Lu, 2018).

To address this limitation, we sought to identify cellular factors that limit SCRIBE’s DNA writing efficiency. We reasoned that cellular factors that reduce the stability of intracellular single-stranded DNA (ssDNA) and those that limit the incorporation of introduced lesions are likely to be involved in limiting the efficiency of this retroelement-mediated DNA writing system. We thus systematically knocked out genes encoding the mismatch repair (MMR) and exonucleases that, respectively, are thought to affect recombination efficiency and intracellular stability of ssDNAs (Costantino and Court, 2003; Sawitzke et al., 2011). We measured SCRIBE genome editing efficiency in these different cellular knockout backgrounds in DH5αPRO cells, which overexpress *lacI* and *tetR*, using a *kanR* reversion assay (hereafter referred to as *kanR*_OFF_ cells) (Farzadfard and Lu, 2014). In this assay, two premature stop codons within a genomic *kanR* cassette (*kanR*_OFF_) are reverted back to the wild-type (WT) sequence by recombineering of intracellularly expressed ssDNAs (ssDNA(*kanR*)_ON_). The retron cassette, which expresses ssDNA(*kanR*)_ON_, as well as the Beta protein, which promotes ssDNA-mediated recombination, were placed in a synthetic operon (dubbed SCRIBE(*kanR*)_ON_) under the control of an isopropyl β-D-1-thiogalactopyranoside (IPTG)-inducible promoter and expressed from a plasmid (Fig. 2A) (Farzadfard and Lu, 2014). The SCRIBE writing efficiency in cells harboring the SCRIBE(*kanR*)_ON_ plasmid was assessed by measuring the recombinant frequency [the ratio of kanamycin-resistant (Kan^R^) cells to total viable cells] in the population in the presence or absence of IPTG induction.

**Figure 2.**
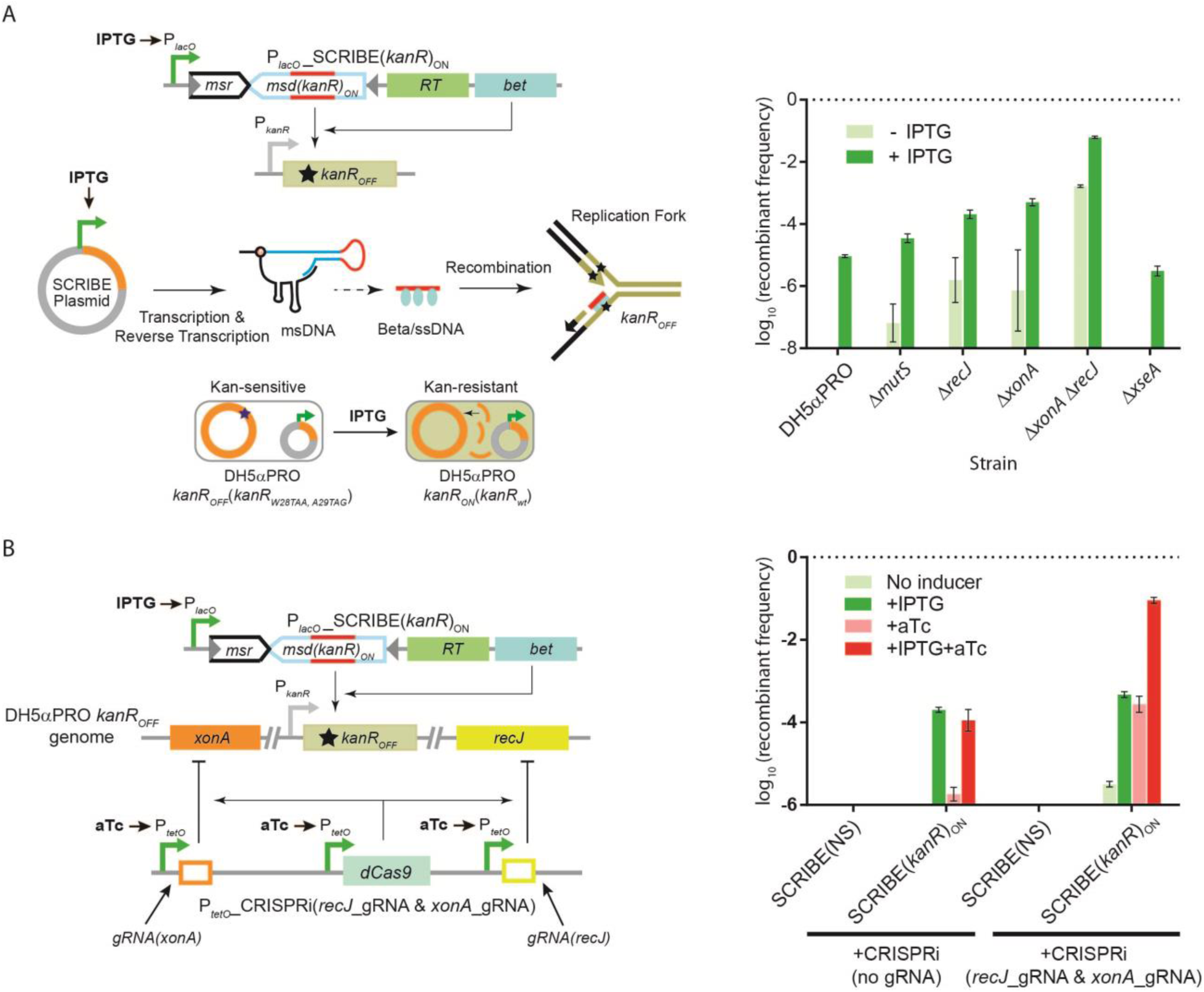
Optimizing SCRIBE DNA Writing Efficiency. **(A)** SCRIBE DNA writing efficiency in different knockout backgrounds in *E. coli* DH5αPRO determined by a *kanR* reversion assay (see Methods). DNA writing efficiency in the Δ*xonA* Δ*recJ* was increased >10^4^-fold relative to the wild-type background. Error bars indicate standard errors for three biological replicates. **(B)** Combining IPTG-inducible SCRIBE and aTc-inducible CRISPRi system (to knockdown cellular exonucleases (*xonA* and *recJ*)) in the WT DH5αPRO strain enables efficient DNA memory recording and dynamic genome engineering with reduced background. Error bars indicate standard errors for three biological replicates.

As shown in Fig. 2A, deactivating the MMR system (Δ*mutS*) resulted in only a modest increase in recombination efficiency. This slight increase may reflect the fact that mismatches longer than three base pairs are poorly recognized by the MMR system (Sawitzke et al., 2011). Knocking out *xseA*, an ssDNA-specific exonuclease that converts large ssDNA substrates into smaller oligonucleotides (Chase and Richardson, 1974), slightly reduced recombination efficiency. On the other hand, knocking out either *recJ* or *xonA*, which respectively encode 5’- and 3’-specific ssDNA exonucleases, significantly increased the recombinant frequency, suggesting that SCRIBE performance is limited by the availability of intracellular recombinogenic ssDNAs (see Supplementary Materials and Fig. S1). Knocking out both *recJ* and *xonA* simultaneously increased the recombinant frequency even further, resulting in a >10^4^-fold increase over the wild-type background. The editing efficiency of this improved DNA writing system is comparable with the highest efficiencies reported for oligo-mediated recombineering (∼10%) (Pines et al., 2015; Sawitzke et al., 2011). Furthermore, consistent with oligo-mediated recombineering, we found the optimum length of homology arm between HiSCRIBE-generated ssDNA template and its target to be ∼35 bps (Fig. S1).

Knocking out cellular exonucleases also increased the background recombination rate to some extent (Fig. 2A), which we speculate is likely due to recombination of ssDNA intermediates generated by the degradation of the template plasmid that persists in cells in the absence of cellular exonucleases (see Supplementary Materials and Fig. S1). To reduce the basal activity of the DNA writer, and to demonstrate that high-efficiency DNA recording is not limited to a specific genetic background, rather than using a knockout background, we conditionally knocked-down *recJ* and *xonA* exonucleases in the WT background using CRISPR interference (CRISPRi) (Qi et al., 2013). We cloned dCas9 and guide RNAs (gRNAs) targeting these two exonucleases under the control of anhydrotetracycline (aTc)-inducible promoters (Fig. 2B). We then co-transformed the DH5αPRO *kanR*_OFF_ reporter strain with this plasmid along with the IPTG-inducible SCRIBE(*kanR*)_ON_ plasmid. Induction of either the SCRIBE or CRISPRi system resulted in a modest increase in the recombination efficiency, while co-induction of both systems resulted in an increase in recombination efficiency of >10^4^-fold over uninduced cells (Fig. 2B). The recombinant frequency was significantly reduced when cells were transformed with a CRISPRi system lacking the gRNAs. No recombinants were detected in cells that were transformed with a non-targeting SCRIBE(NS) plasmid. These results further support that cellular exonucleases limit SCRIBE genome editing efficiency and demonstrate that efficient DNA recording can be performed in genomically unmodified cells by combining SCRIBE and CRISPRi, with significantly less background compared to the exonuclease knockout strain. This feature could enable building more robust DNA recorders and other computing-and-memory circuits that use DNA as the computing substrate, without the need to genetically engineer target cells beforehand.

In addition to molecular recording applications, such as linking a transcriptional signal to a precise mutation in the genome, HiSCRIBE DNA writers can be used for genome editing applications for which maximal DNA writing efficiencies are desired. Oligo-mediated recombineering can introduce desired modifications into a bacterial genome, but in this technique, synthetic oligos are introduced to target cells transiently (via electroporation) and have a very short intracellular half-life. Due to these shortcomings and the simultaneous presence of multiple replication forks in bacteria, the theoretical editing efficiency of oligo-mediated recombineering is limited to 25%, while the practical editing efficiency is often limited to a few percents (Pines et al., 2015; Sawitzke et al., 2011). Thus, multiple rounds of recombineering are needed to improve efficiency and additional screening steps are required to obtain desired mutants. In addition, to achieve such efficiencies, it is often necessary to modify the host by knocking out the MMR system, which, in turn, could elevate the global mutation rate and leads to undesirable genome-wide off-target mutations (Schaaper and Dunn, 1987). In contrast, HiSCRIBE provides a persistent intracellular source of recombinogenic oligos over many generations and can be introduced to cells even with low-efficiency delivery methods, thus bypassing the above-mentioned limitations.

To demonstrate high-efficiency genome editing by HiSCRIBE writers, we created a gene editing-optimized HiSCRIBE system by engineering a stronger Ribosome Binding Site (RBS) to overexpress Beta (Fig. S2). Using this system, we sought to change two consecutive leucine codons in the *galK* ORF in the MG1655 Δ*recJ* Δ*xonA* strain (hereafter referred to as MG1655 *exo^-^* strain) to synonymous codons. Cells were transformed with the HiSCRIBE(*galK*)_SYN_ plasmid, which encodes an ssDNA with mismatches in three nucleotides to *galK* in order to write synonymous leucine codons into *galK* while effectively avoiding the MMR system (Sawitzke et al., 2011). We plated these cells on agar and then monitored the conversion of the genomic *galK*_WT_ allele to the *galK*_SYN_ allele in transformants at 24 hours after transformation (∼30 generations) by colony PCR followed by Sanger sequencing as well as Illumina sequencing. As shown in Fig. 3A (middle panel), Sanger chromatograms obtained from colonies at this stage showed mixed peaks at the targeted nucleotides, indicating the presence of both *galK*_WT_ and *galK*_SYN_ alleles within single colonies. Sequencing these amplicons using Illumina MiSeq indicated that ∼60% of the *galK*_WT_ allele was converted to *galK*_SYN_ after one day (Fig. 3A, bottom panel). Since Beta-mediated recombineering is a replication-dependent process (Farzadfard and Lu, 2014; Huen et al., 2006), the frequency of recombinants in HiSCRIBE-expressing populations should increase with more generations. We thus re-streaked the transformants on new plates to allow additional time for writing. After an additional day of growth, Sanger sequencing of *galK* PCR amplicons from these new colonies revealed that the conversion of the *galK*_WT_ allele to *galK*_SYN_ was so efficient that the *galK*_WT_ allele was below the limit of detection (Fig. 3A). Illumina sequencing of the amplicons further confirmed that ∼100% of *galK*_WT_ allele within individual colonies was converted to *galK*_SYN_. When cells were transformed with a non-specific HiSCRIBE(NS) plasmid, no modified alleles were detected by sequencing.

**Figure 3.**
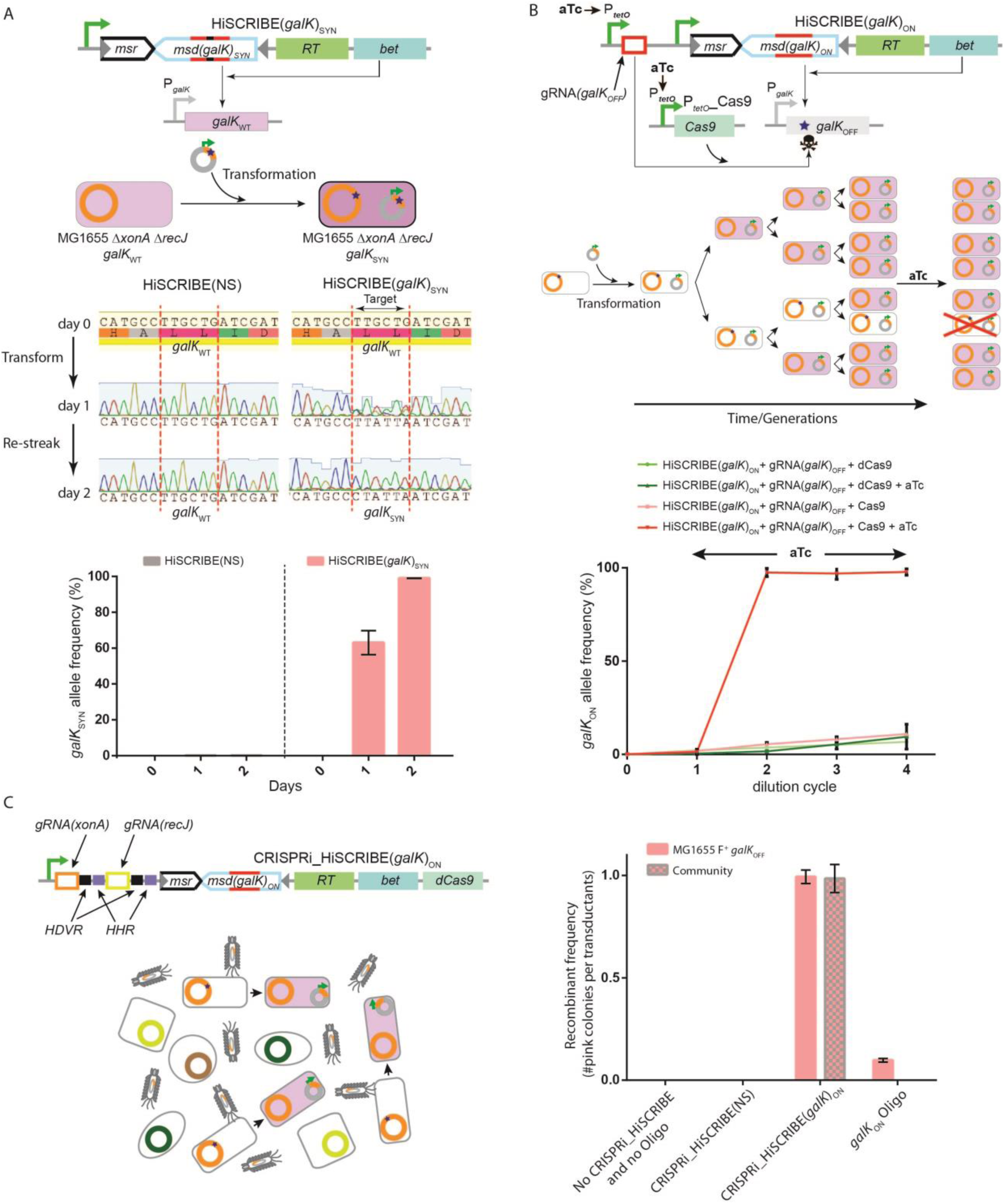
Highly-efficient, specific, scarless, and *cis*-element-independent editing of bacterial genomes by HiSCRIBE in clonal populations and within synthetic communities. **(A)** HiSCRIBE enables highly efficient genome editing in clonal populations. HiSCRIBE was used to convert two consecutive leucine codons in the *galK* locus of MG1655 *exo^-^* cells to synonymous codons. Cells were transformed with the HiSCRIBE(*galK*)_SYN_ plasmid and the conversion of the *galK*_WT_ to *galK*_SYN_ was monitored 24 hours after transformation by PCR amplification of the *galK* locus of the transformants followed by Sanger sequencing (middle panel) as well as Illumina sequencing (bottom panel). Re-streaking the transformants on new plates and growing cells for an additional 24 hours led to the ∼100% conversion of the *galK*_WT_ allele to *galK*_SYN_ in all the tested transformants (also see Supplementary Materials and Fig. S3). No allele conversion was observed in cells that had been transformed with the non-specific HiSCRIBE(NS) plasmid. **(B)** Combining HiSCRIBE DNA writing with aTc-inducible CRISPR-Cas9 nuclease-mediated counterselection of unedited wild-type alleles increases the rate of enrichment of modified alleles within MG1655 *galK*_OFF_ *E. coli* population (see Methods). Error bars indicate standard deviation for three biological replicates. **(C)** Genome editing within a bacterial community via phagemid-mediated delivery of the HiSCRIBE system. Target cells (*E. coli* MG1655 *galK*_OFF_ F^+^ Str^R^) either as a clonal bacterial population or mixed with a stool-derived bacterial community were incubated with HiSCRIBE(*galK*_ON_) or HiSCRIBE(NS) phagemid particles and DNA writing efficiency in the *galK* locus was assessed by the *galK* reversion assay (see Methods). Recombinant frequency was calculated as the ratio of pink (galactose fermenting) colonies to target cell transductants. As additional controls, we used oligo-mediated recombineering with a synthetic *galK*_ON_ oligo to edit reporter cells harboring a recombineering pKD46 plasmid either as a clonal population or in the context of a bacterial community. Recombinant frequency was calculated as the ratio of pink (galactose fermenting) colonies to total viable reporter cells. Transduction efficiencies of the HiSCRIBE phagemids are presented in Fig. S4. Error bars indicate standard errors for three biological replicates.

We further assessed the DNA writing frequency in the entire population using a screenable plating assay and observed that more than 99% of transformants [colony forming units (CFUs)] in the population underwent intended DNA editing after receiving the HiSCRIBE plasmid (see Supplementary Materials and Fig. S3). As in the previous experiment, more than 99% of WT alleles within each CFU were converted into mutated alleles within 2 days (∼60 generations). Overall, these results demonstrate that HiSCRIBE is a highly efficient, broadly applicable, and scarless genome writing platform that can achieve ∼100% editing efficiency at both the single-cell and population level without requiring any *cis*-encoded sequence on the target, double-strand DNA breaks, or selection.

## Increasing the Rate of Allele Enrichment by Nucleotide-Resolution Counter-selection Using CRISPR-Cas9 Nuclease

The enrichment of a mutant allele within a population directly correlates with its fitness. In the absence of a selective advantage, it may take many generations for a neutral allele to accumulate within a population (Farzadfard and Lu, 2014). As demonstrated in Fig. 3A, the editing-optimized HiSCRIBE by itself can achieve ∼100% editing efficiency over the course of two days (∼60 generations) during which the desired mutation accumulates in a replication-dependent fashion. We sought to increase the rate of this process by putting a selective pressure against the WT allele at the nucleotide level using CRISPR-Cas9 nuclease. To this end, we first constructed a *galK*_OFF_ reporter strain by introducing two premature stop codons into the MG1655PRO Δ*recJ* Δ*xonA* strain (hereafter referred to as MG1655PRO *exo^-^ galK*_OFF_ reporter strain). We encoded an aTc-inducible gRNA against the *galK*_OFF_ allele into the HiSCRIBE(*galK*)_ON_ plasmid, which expresses ssDNA with the same sequence as the WT *galK*. This plasmid was then transformed into MG1655PRO *exo*^-^ *galK*_OFF_ reporter cells containing either aTc-inducible Cas9 or dCas9 (as a negative control) plasmids. Single colonies of transformants were grown, diluted, and regrown for multiple cycles in the presence or absence of aTc. The dynamics of *galK* alleles frequency in different cultures were monitored throughout the experiment by PCR amplification and deep-sequencing of the *galK* locus at different time points. As shown in Fig. 3B, *galK*_ON_ allele was enriched in all the cultures over time, further confirming that genome editing via HiSCRIBE is a replication-dependent process. However, upon induction with aTc, the *galK*_ON_ alleles were quickly enriched in cells expressing Cas9 compared to cells expressing dCas9 and comprised ∼99% of *galK* alleles 12 hours after induction. These results demonstrate CRISPR-Cas9 nuclease activity, which is deleterious by itself if targeted against a bacterium’s own genome (Caliando and Voigt, 2015; Citorik et al., 2014), can be combined with HiSCRIBE genome writing to induce selective sweeps and accelerate the enrichment of desired alleles in a population.

## Efficient and Specific Genome Editing of Bacteria within Communities

Traditional recombineering techniques rely on high-efficiency delivery methods, such as electroporation or natural competence, for the introduction of synthetic oligos to the cells. This reliance, however, limits the applicability of these techniques to certain laboratory conditions (e.g., highly electrocompetent cells grown to mid-log phase in test tubes). Unlike these traditional techniques, which cannot be applied to edit bacterial genome within complex communities or *in situ*, HiSCRIBE can be encoded on plasmids and delivered to cells via low transformation efficiency methods, such as chemical transformation, or via transduction or conjugation (Fig. S4), thus greatly expanding the applicability of recombineering techniques to complex bacterial communities and intractable bacteria.

To demonstrate the possibility of delivering HiSCRIBE by these alternative methods, we first encoded HiSCRIBE on an M13 phagemid and used it to target and edit specific cells within a synthetic bacterial community. We introduced our target strain, *E. coli* MG1655 *galK*_OFF_ F^+^ Str^R^ (which encodes the receptor for M13 bacteriophage on F plasmid), into an undefined bacterial community derived from mouse stool at a 1:100 ratio to make a synthetic bacterial community. To reduce the number of plasmids that needed to be delivered into target cells, we placed both HiSCRIBE(*galK*)_ON_ and the CRISPRi cassette targeting *recJ* and *xonA* exonucleases in a single synthetic operon, referred to as the CRISPRi_HiSCRIBE(*galK*)_ON_ operon (Fig. 3C). To allow for *in vivo* processing and release of these gRNAs from the synthetic operon transcripts, gRNAs were flanked by a Hammerhead Ribozyme (*HHR*) and a hepatitis delta virus Ribozyme (*HDVR*) (Gao and Zhao, 2014). We cloned this synthetic operon into a plasmid harboring the M13 bacteriophage packaging signal. CRISPRi_HiSCRIBE-encoding M13 phagemid particles were produced in an M13 packaging strain, purified, and added to the synthetic community or the reporter strain alone. The target cells were then scored on MacConkey + Streptomycin (Str) + galactose (gal) plates for the ability to metabolize galactose, indicated by pink coloring (*galK* reversion assay, see Methods).

As shown in Fig. 3C, more than 99% of the reporter cells that received CRISPRi_HiSCRIBE(*galK*)_ON_ phagemids formed pink colonies on the indicator plates, demonstrating successful editing of targeted cells within a complex community. No pink colonies were observed in the negative control, in which the bacterial community was transduced with non-specific CRISPRi_HiSCRIBE(NS) phagemid particles. As an additional control, we introduced *galK*_ON_ oligo into reporter cells harboring the pKD46 recombineering plasmid, either in a clonal population or within the synthetic community, using an established recombineering protocol (Datsenko and Wanner, 2000; Sawitzke et al., 2011). Consistent with previous reports, we observed ∼10% recombineering efficiencies in the clonal population of the reporter strain. However, no recombinant pink colonies were obtained when reporter cells were contained within the synthetic community, further confirming that highly efficient delivery of oligos, as needed for traditional recombineering, is not achievable in bacterial communities. We further showed that conjugation, a common strategy for horizontal gene transfer in natural bacterial communities, can be used to deliver the HiSCRIBE plasmid for genome editing within bacterial communities (Fig. S4B). These results demonstrate that diverse strategies can be used to deliver HiSCRIBE constructs into complex bacterial communities with the potential for *in situ* genome-editing applications.

## Recording Spatial Information into DNA Memory

A useful feature of the HiSCRIBE system is that, unlike oligo-mediated recombineering, high-efficiency DNA writing can be linked to and triggered by biological processes. This feature could be especially useful to study events and interactions that occur in biological systems, such as cell-cell interactions within bacterial communities and biofilms, that are transient and thus hard to study in high throughput or with high resolution, especially in their native contexts. Enabled by the highly-efficient HiSCRIBE DNA writers, we devised a barcode joining strategy to uniquely mark and permanently record such transient interactions into DNA. The recorded memory can be retrieved via high-throughput sequencing to map and study the interactome with high resolution and throughput, even after samples and interactions are disrupted.

We sought to demonstrate this concept by mapping conjugation events between bacterial populations. To this end, we first designated two neighboring 6 bp sequences on the *galK* locus as memory registers. We then constructed a series of HiSCRIBE(Reg1)_r-barcode_ and HiSCRIBE(Reg2)_d-barcode_ plasmids, each encoding a different barcoded ssDNA template. These plasmids each write a unique 7 bp DNA sequence (1 bp writing control + 6 bp barcode) on the first and the second registers, respectively (Fig. 4A). The writing control nucleotide was designed as a mismatch to the unedited register and used to selectively amplify edited registers (see Methods). We introduced the HiSCRIBE(Reg1)_r-barcode_ plasmids into the MG1655 *exo^-^* strain to make a set of conjugation recipient populations. Upon transformation, these plasmids wrote a unique barcode in the first genomic register in these cells (Register 1), and uniquely marked these recipient populations. HiSCRIBE(Reg2)_d-barcode_ plasmids, harboring an RP4 origin of transfer, were used to transform MFDpirPRO cells to make a set of conjugation donor populations. Upon transfer from donor to recipient, these plasmids write a unique barcode in Register 2 in recipient cells. Sequencing the consecutive Register 1 and Register 2 in recipient genomes yielded a record of this interaction (Fig. 4A).

**Figure 4.**
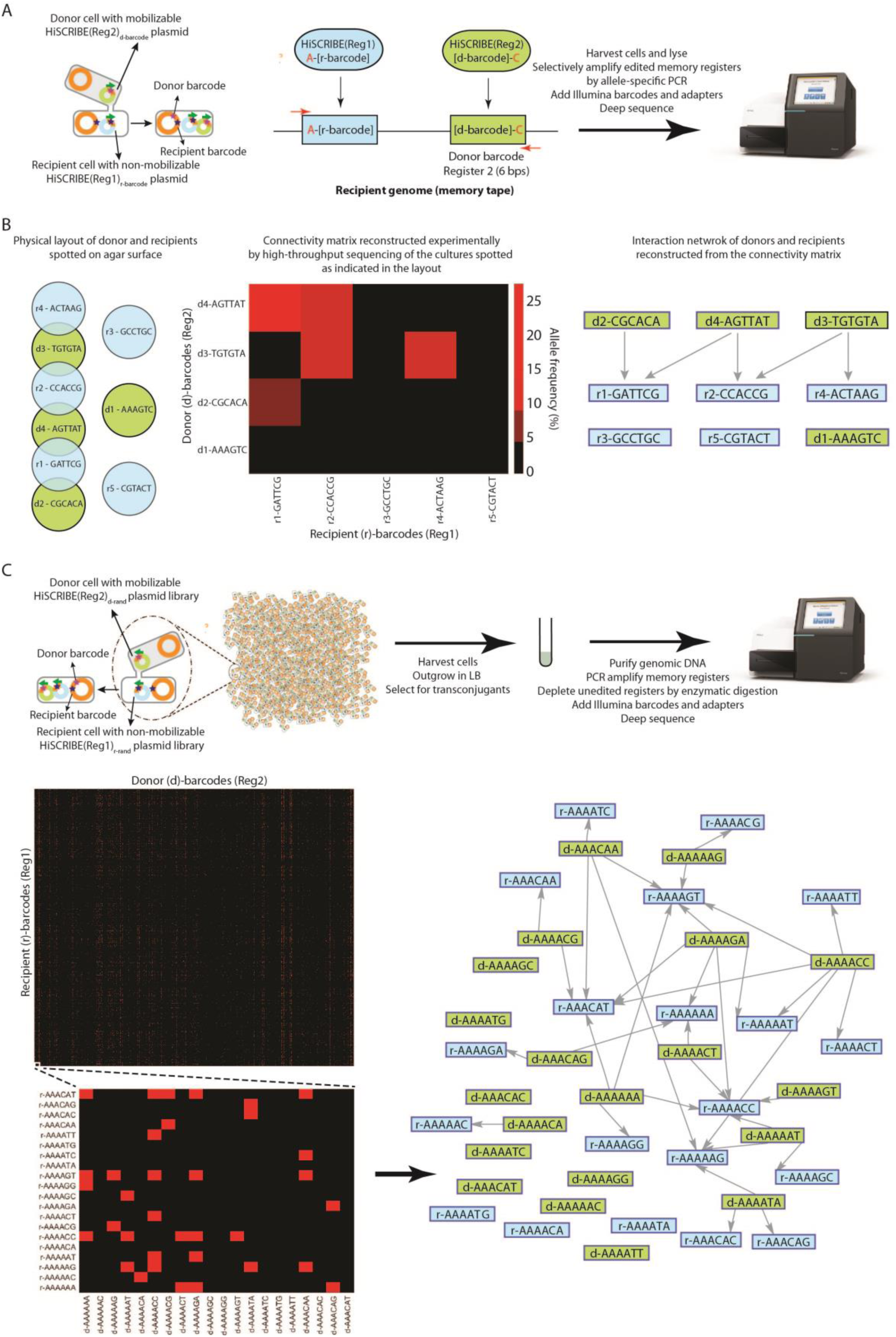
Mapping spatial patterns and the connectome of conjugative mating pairs in bacterial populations. **(A)** Schematic representation of the barcode joining strategy used to record pairwise interactions (conjugation events) between conjugative pairs of bacteria using HiSCRIBE-based DNA writing. Upon successful conjugation, the interactions between a recipient cell and donor cell are recorded into neighboring DNA memory registers in the recipient cell genome. The edited registers are then amplified using allele-specific PCR (to deplete non-edited registers) and the identities of the interacting partners are retrieved by sequencing. The nucleotide shown in red in each register represents a single nucleotide that was included in each barcode to distinguish between unedited and edited registers. These “writing control” nucleotides were then used to selectively amplify edited registers by allele-specific PCR using primers that match these nucleotides but not to unedited registers. **(B)** Detecting the spatial organization of clonal bacterial populations. Clonal populations of donors and recipients harboring HiSCRIBE-encoded “d-*barcode*” (green circles) and “r-*barcode*” (blue circles), respectively, were spotted on nitrocellulose filters that were then placed on the agar surface in the patterns shown in the left panel. Conjugation mixtures were harvested and the memory registers were amplified by allele-specific PCR and sequenced by Illumina sequencing (see Methods). Recorded barcodes in the two consecutive memory registers were parsed and the donor-recipient population connectivity matrix was calculated based on the percentage of reads corresponding to each possible pair-wise interaction of donors and recipient barcodes. The heatmap representation of the retrieved connectivity matrix (middle panel), as well as the corresponding interaction network (right panel), are shown. Red boxes in the heatmap depict connected barcodes, indicating that a conjugation event from the corresponding donor population resulted in HiSCRIBE transfer and subsequent recording of the donor barcode into the specific recipient genome. In the interaction network, donor and recipient barcodes are indicated by green (“d-*barcode*”) and blue (“r-*barcode*”) rectangles, respectively. Data obtained from additional spatial patterns for bacterial populations are provided in Fig. S5. **(C)** The strategy used to map conjugation events between individual pairs of donor and recipient cells as a proxy for a conjugative connectome using randomized HiSCRIBE libraries. The connectivity matrix was obtained using the method described in (B). Due to the large size of the connectivity matrix (∼16 million elements), a submatrix for the first 20 (alphabetically sorted) barcodes of donors and recipients in one of the samples is shown in the inset. The y- and x-axis show recipient genomic barcodes (recorded in Register 1) and donor barcodes (recorded in Register 2), respectively. The corresponding interaction subnetwork for the presented connectivity submatrix is shown on the right. Entire connectivity matrices for three parallel conjugation experiments are provided in Supplementary File 1.

Using this barcode joining strategy, we first demonstrated that the interaction between a barcoded donor population and a barcoded recipient population could be successfully recorded and faithfully retrieved by allele-specific PCR of conjugation mixtures followed by Sanger sequencing (Fig. S5). To this end, we spotted a donor population with a single donor barcode on filter paper, overlapped it with another filter paper with a recipient population containing a single recipient barcode, and then confirmed that our retrieval process was correct (Fig. S5A). We then constructed more complex spatial layouts by overlapping multiple different barcoded donor populations and barcoded recipient populations. We demonstrated that allele-specific PCR combined with high-throughput sequencing could faithfully retrieve conjugative interactions between the distinct barcoded donor and recipient populations laid down in different patterns (Fig. 4B and Fig. S5B).

After validating that the barcode joining strategy can be used to map interactions between barcoded bacterial populations, we next sought to map cell-cell interactions at the single-cell resolution as an example of a “cellular connectome”. In this experiment, we used donor and recipient populations harboring pooled randomized barcodes that uniquely barcode individual cells in each population. Specifically, we constructed a pooled recipient population, harboring a HiSCRIBE(Reg1)_r-rand_ plasmid library that encoded an ssDNA library with 6 randomized nucleotides targeting Register 1 in the *galK* locus. We also created a pooled donor library by transforming MFDpirPRO cells with a conjugative HiSCRIBE(Reg2)_d-rand_ plasmid library that similarly encoded an ssDNA library with 6 randomized nucleotides targeting Register 1 in the *galK* locus. To test this method of recording mating interactions at the single-cell level, donor and recipient populations were mixed and spotted on filter paper on a solid agar surface to allow for conjugation of the HiSCRIBE(Reg2)_d-rand_ library from donors to recipients (Fig. 4C). Samples were then disrupted and grown in liquid cultures to allow for propagation and amplification of rare conjugated alleles. The two neighboring DNA memory registers were amplified as a single amplicon by PCR, enzymatically digested to remove non-edited registers that contained parental restriction sites, and deep sequenced (Fig. S6A, see Methods). Connectivity matrices between members of donor and recipient populations were then deduced based on the DNA barcodes obtained in the two specified memory registers (Fig. 4C and Supplementary File 1). Unique variants in the HiSCRIBE-targeted registers were three orders of magnitude higher than in randomly chosen non-targeted regions (Fig. S6C), indicating successful recording of conjugation events.

To better understand the conjugation events between different population members, we analyzed the frequencies of interacting donor and recipient barcodes in three parallel experiments. The degree of donor barcodes, which was defined as the number of different connections that each unique donor barcode makes with recipient barcodes, was well correlated among the three parallel experiments (Fig. S7). This suggests that in these conjugation mixtures, the number of conjugation events in which each donor barcode participates is independent of the identity of their interacting partners (i.e., the recipient barcodes) and likely depends on the rate of transfer of donor barcodes, which itself is likely to be a function of the frequency of these barcodes in the population and the efficiency of transfer of each individual barcode. On the other hand, we observed a weak correlation between the degree of recipient barcodes, which was defined as the number of different connections that each unique recipient barcode makes with donor barcodes, in the three parallel experiments. This indicates that the number of donor barcodes that interact with each unique recipient barcode is different in each sample and suggests that other factors, such as the identity and frequency of donors in each iteration of conjugation, could affect the rate of successful conjugation and barcode transfer/writing in recipients.

With these proof-of-concept experiments, we demonstrated that an efficient DNA writer that is coupled with biological processes can be used to memorize transient information, such as spatial patterns and cell-cell mating events between bacterial strains, into genomic DNA in their native context. This strategy allows reducing multi-dimensional interactome space to one-dimensional DNA sequence space which can then be later retrieved by sequencing. We call this strategy “DNA imaging” as it is conceptually analogous to the traditional imaging techniques in which spatial information is transformed and permanently captured in a recording medium. With the described strategy, using two 6-nucleotide memory registers, up to 4^12^ ∼ 16 million unique interactions can be theoretically recorded. The recording capacity can be increased by using larger barcodes. In our experiment, we could detect ∼1% of the theoretical recording capacity (Fig. S6C), although increasing the sequencing depth or conjugation sample size could help to increase barcode recovery. While only pairwise interactions were recorded in this proof-of-concept experiment, in principle, multiple interactions can be recorded into adjacent DNA registers to map multidimensional interactomes with high-throughput sequencing, particularly as sequencing fidelity and read lengths continue to improve. We envision that DNA imaging by HiSCRIBE and other analogous efficient and precise DNA writing systems could be used for high-throughput and high-resolution mapping of cellular organizations and connectomes, as well as other types of intracellular transient interactions, in complex and opaque environments where traditional imaging techniques are not applicable.

## Continuous *in vivo* Evolution

Evolution is a continuous process of genetic diversification and phenotypic selection that tunes the genetic makeup of living organisms and maximizes their fitness in a given environment over evolutionary timescales. Evolutionary design is a powerful approach for engineering living systems. However, in many cases, natural mutation rates are not high enough to allow desirable genetic changes to be accessible on practical timescales. Efficient and *cis*-element-independent HiSCRIBE DNA writers could enable the targeted diversification of desired loci *in vivo* in a continuous and temporally and spatially programmable manner. Targeted diversity generation could be coupled with a continuous selection or screening setup to achieve adaptive writing and tune cellular fitness continuously and autonomously with minimal human intervention (Fig. 5A).

**Figure 5.**
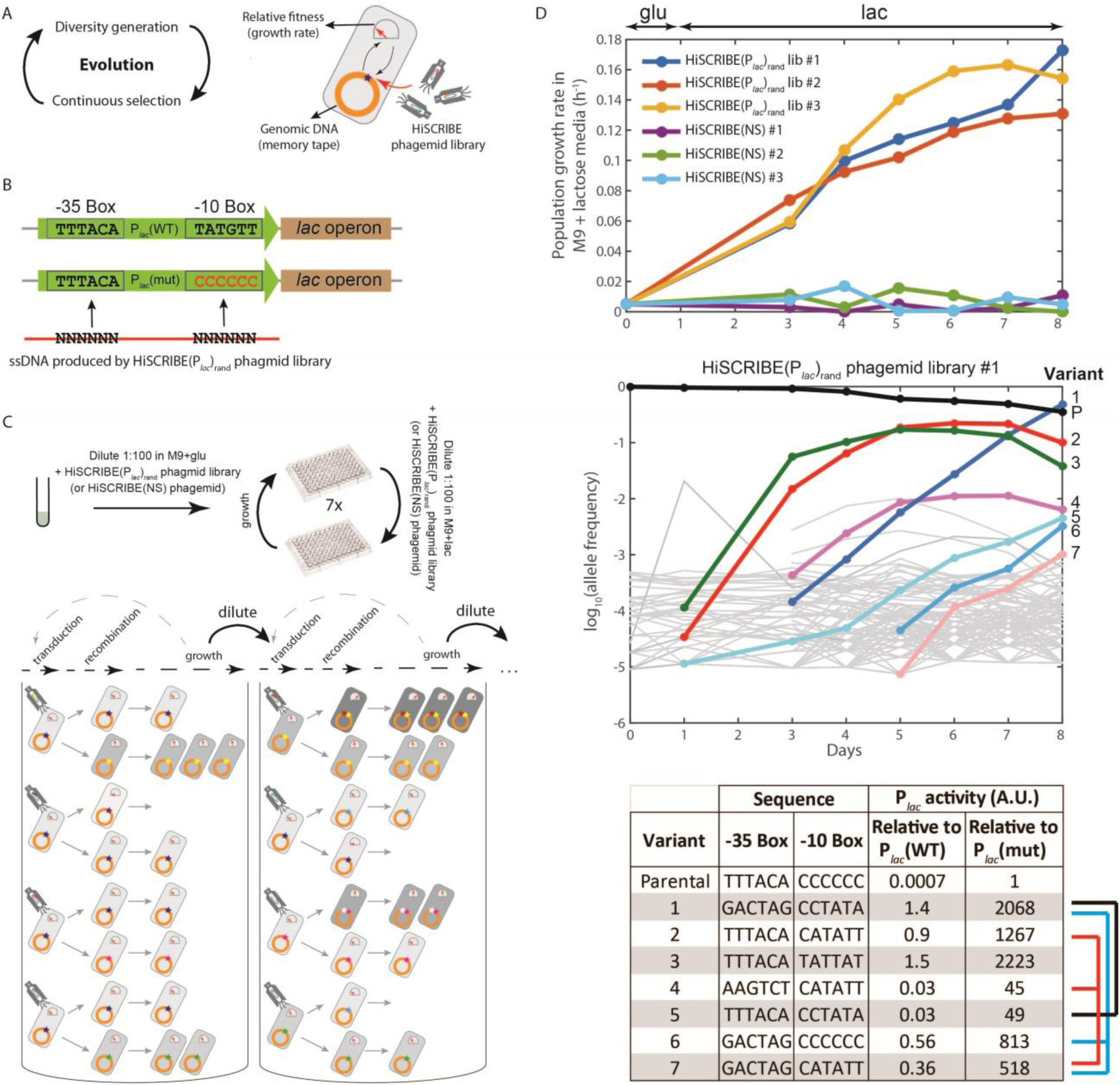
Continuous evolution of a desired genomic locus via HiSCRIBE. **(A)** diversity generation enabled by HiSCRIBE can be coupled with continuous selection to accelerate the rate of evolution of desired target sites. A randomized HiSCRIBE library was encoded on phagemids that were continuously delivered into cells. In the presence of a selective pressure, HiSCRIBE-mediated mutations lead to adaptive genetic changes that increase fitness. An increase in fitness results in faster replication and amplification of the associated genotype, increasing the chance that cells containing the genotype can undergo additional rounds of diversification. **(B)** The sequences of -35 and -10 boxes of the wild-type P*_lac_* (P*_lac_*(WT)) and mutated P*_lac_* (P*_lac_*(mut)) targeted by a phagemid-encoded randomized HiSCRIBE(P*_lac_*)_rand_ library in the evolution experiment. **(C)** Schematic representation of the evolution experiment. The -35 and -10 boxes of the P*_lac_* locus were targeted with an ssDNA library produced *in vivo* from a HiSCRIBE phagemid library delivered by transduction. Cells that acquired beneficial mutations in their P*_lac_* locus were expected to metabolize lactose better (indicated by darker gray shading) and be enriched in the population over time. **(D)** Growth rate profiles of cell populations exposed to HiSCRIBE(P*_lac_*)_rand_ and HiSCRIBE(NS) (top) as well as the dynamics of P*_lac_* alleles over the course of the experiment are shown as time series for cells exposed HiSCRIBE(P*_lac_*)_rand_ phagemid library (middle). The bottom panel shows the identities of the most frequent alleles at the end of the experiment as well as the fold-change in the β-galactosidase activity of those alleles in comparison to the WT and parental alleles. Alleles that are likely ancestors/descendants are linked by brackets. The dynamics of allele enrichment for cells exposed to HiSCRIBE(NS) and additional parallel evolution experiments are presented in Fig. S8.

To demonstrate this concept, we linked cellular fitness (i.e., growth rate) to a cell’s ability to consume lactose as the sole carbon source. To enable a wide dynamic range in fitness to be explored, we first weakened the activity of the native *lac* operon promoter (P*_lac_*) by introducing mutations into its -10 box (P*_lac_*(mut), Fig. 5B) in *E. coli* MG1655 *exo^-^* strain. Cells with the P*_lac_*(mut) promoter (hereafter referred to as the parental strain) grew poorly in minimal media (M9) when lactose was present as the sole carbon source. We then used a randomized HiSCRIBE phagemid library (HiSCRIBE(P*_lac_*)_rand_) to continuously introduce diversity into the -10 and -35 sequences of this promoter (Fig. 5B, see Supplementary Materials). Starting from an overnight culture, parental cells were diluted into M9 + glucose media and divided into two groups, which were then treated with phagemid particles from either HiSCRIBE(P*_lac_*)_rand_ library or HiSCRIBE(NS). After this initial growth in glucose, cells were diluted and regrown in M9 + lactose in the presence of phagemid particles for six additional rounds to allow for concomitant diversification, selection, and propagation of beneficial mutations (Fig. 5C, see Supplementary Methods). As shown in Fig. 5D (top panel), the overall growth rates of cell populations in lactose increased when they were transduced with the HiSCRIBE(P*_lac_*)_rand_ phagemid library. In contrast, the growth rates of cell populations exposed to the control HiSCRIBE(NS) phagemid particles did not change over time. These results demonstrate that the randomized HiSCRIBE library can introduce targeted diversity into desired loci (-10 and -35 boxes of the P*_lac_* promoter) that leads to increase in the fitness of the population under selection over relatively short timescales, much faster than what can be achieved by natural Darwinian evolution (i.e., in cells transformed with non-targeting HiSCRIBE(NS)).

To monitor the dynamics of mutant allele enrichment in these cultures, at different time points over the course of the experiment, the P*_lac_* region was PCR-amplified and deep-sequenced. The diversity and frequency of P*_lac_* alleles in samples that had been exposed to the HiSCRIBE(NS) phagemid did not change significantly over time and the parental allele comprised ∼100% of the population at all analyzed time points (Fig. S8A and B). Further inspection of the rare variants observed in these samples revealed mostly single nucleotide changes compared to the parental allele, suggesting that these arose from sequencing errors. On the other hand, the diversity of P*_lac_* alleles greatly increased in cultures that were exposed to the HiSCRIBE(P*_lac_*)_rand_ phagemid library when they were initially grown in the M9 + glucose condition (Fig. S8A). This initial increase in allele diversity was followed by a significant drop upon dilution of the cells in lactose media, likely due to sampling drift and strong selection for alleles that allow for lactose metabolism. Throughout the experiment, however, the number of unique variants remained significantly higher in the HiSCRIBE(P*_lac_*)_rand_ cultures than in the negative controls. Moreover, the frequency of P*_lac_* alleles from samples that had been exposed to HiSCRIBE(P*_lac_*)_rand_ changed dynamically over time (Fig. 5D, middle panel). Notably, by the end of the experiment, the frequency of the parental allele dropped to less than 50% and one variant (variant #1) became the dominant allele in the population. Further analysis of the frequent variants within the diversified population indicated that multiple mutations occurred in the -10 and -35 boxes in discrete steps, in which secondary mutations arising on top of primary mutations led to an increase in fitness (Fig. 5D, bottom panel). For example, based on allele enrichment and P*_lac_* activity data (see below), the dominant allele (variant #1) was likely produced from an initial, less active mutant (variant #5) and subsequently took over the population due to its higher fitness (i.e., P*_lac_* activity). The sequences of the enriched variants that evolved in this experiment were especially AT-rich (Fig. 5D, bottom panel, and Fig. S8C), as is expected from the canonical sequences of these regulatory elements in *E. coli*.

To validate that the enriched variants were indeed responsible for the observed increase in fitness, we reconstructed these variants in the parental strain background and assessed their activity by measuring β-galactosidase activity. As shown in Fig. 5D (bottom panel), all of these variants showed a significant increase in β-galactosidase activity over the parental variant, indicating successful tuning of the activity of the P*_lac_* promoter. For example, the dominant variant at the end of the experiment (variant #1) exhibited a >2000-fold increase in β-galactosidase activity relative to the parental strain, corresponding to a 1.4-fold increase over the wild-type P*_lac_* promoter.

These results demonstrate that once coupled to a continuous selection or screening (Rogers et al., 2016), HiSCRIBE can be used for continuous and autonomous diversity generation in desired target loci, thus enabling easy and flexible continuous evolution experiments with minimal requirement for human intervention. In the current setup, the continuous diversity generation system relies on the delivery of phagemid-encoded HiSCRIBE variants that compete for writing on the target locus once inside the cells. In future work, incorporating a conditional origin of replication into phagemids or conjugative plasmids may help to increase the rate of evolution by enforcing writing and curing steps in a more controlled fashion.

## Discussion

Recently, several DNA writing technologies for recording molecular events into the DNA of living cells have been described. Memory recording using site-specific recombinases (Roquet et al., 2016), CRISPR spacer acquisition (Shipman et al., 2016), Cas9 nuclease (Perli et al., 2016), and base editing (Farzadfard et al., 2019) requires *cis*-encoded elements (e.g., recombinase sites, CRISPR repeats, PAM domains, etc.) and thus is confined to certain loci. In contrast, HiSCRIBE writers do not require any *cis*-encoded element on the target and thus open up the entire genome for efficient genome editing and molecular recording applications. Furthermore, HiSCRIBE enables active and dynamic modification of bacterial genomes without generating double-strand DNA breaks, which may help to reduce associated cytotoxicity and unwanted chromosomal rearrangements. This feature is especially important for genome editing in the context of bacterial communities and evolutionary engineering applications, where fitness costs could be deleterious for the targeted population. Additionally, unlike genome editing strategies that rely on counterselection by CRISPR-Cas9 nucleases (Jiang et al., 2013), HiSCRIBE does not require the presence of a PAM sequence on the target and can operate without the need for a counter-selection system, thereby allowing one to perform multiple rounds of allele replacement on the same target, a property which is especially important for evolutionary engineering applications. Furthermore, HiSCRIBE template plasmids can serve as unique barcodes to identify and track mutations and their enrichment in genome-wide trait optimization scenarios, a challenge for traditional recombineering-based approaches (Zeitoun et al., 2015). Additionally, by providing a sustainable source of mutagenic oligos *in vivo*, the HiSCRIBE system, upon further optimization, could help to bypass current limitations in performing recombineering in hard-to-transform hosts in which Beta or its homologs are functional (Corts et al., 2019; Wannier et al., 2020) and expand the applicability of recombineering-based techniques to *in situ* conditions.

To highlight the power of dynamic recording and autonomous genome engineering enabled by an efficient *in vivo* DNA writing system such as HiSCRIBE, we first demonstrated that spatial information such as cellular patterns and cell-cell interactions can be mapped with high resolution and throughput. Efficient DNA writers could be used to study bacterial spatial organization within biofilms, which has been challenging to do with the traditional techniques (Nadell et al., 2016). In future work, HiSCRIBE could be encoded in phages, conjugative plasmids, or other mobile genetic elements and designed to write similar barcodes near identifiable genomic signatures (e.g., 16S rRNA gene) to assess the *in situ* host range of these mobile elements. In addition, efficient and conditional DNA writers could be used to record other types of transient spatiotemporal events, such as protein-protein interactions, into DNA for high-throughput studies. Furthermore, extending this barcode joining approach to multicellular organisms and mammalian cells using analogous high-efficiency DNA writing technologies, may help to record and map cellular interactions such as neural connectomes (using neural viruses that can pass through synapses as barcode carriers) (Glaser et al., 2015; Peikon et al., 2017; Zador et al., 2012). Although in this study we used our system only to record spatial and temporal biological events, in principle, arbitrary information can be encoded and written into the genomic DNA of living cells (Shipman et al., 2017). For example, digital information (e.g., documents, images, videos, etc.) could be encoded into HiSCRIBE ssDNA templates and written across various genomic loci in living cell populations. The recorded memory could then be retrieved by sequencing the genomic memory registers.

To further demonstrate the utility of efficient and precise DNA writing systems, we used HiSCRIBE writers to continuously and autonomously tune a genomic segment (P*_lac_* promoter) and its connected phenotype (ability to metabolize lactose) in *E. coli*. We demonstrated that HiSCRIBE phagemid libraries and cells comprise a self-contained and rapidly evolving synthetic ecosystem that can continuously and autonomously traverse evolutionary paths imposed by the diversity of the HiSCRIBE library and the applied selective pressure. This platform could facilitate gene resurrection studies, which have so far been limited because of the lack of suitable tools for continuous and targeted *in vivo* mutagenesis. Such studies could provide new insights into accessible evolutionary trajectories (Jermann et al., 1995; Pal et al., 2014; Risso et al., 2013; Thornton, 2004; Weinreich et al., 2006) and empower advanced evolutionary genome engineering approaches. In addition to phagemid delivery, inducible writing (as shown in Fig. 2B) or conjugative delivery of HiSCRIBE libraries (as shown in Fig. 4C) could be linked to selection or screening strategies to enable temporally or spatially restricted diversification and continuous evolutionary engineering of cellular phenotypes. Unlike recombineering-based targeted mutagenesis strategies like Multiplexed Automated Genome Engineering (MAGE), where the library size is limited by the capacity to electroporate synthetic oligos into a limited number of cells, HiSCRIBE diversity generation can be readily scaled-up using alternative delivery methods such as transduction and conjugation. This feature could greatly expand the practical diversity that can be experimentally introduced into a population and the breadth of organisms that can be targeted. Furthermore, unlike MAGE, the diversity generation step for HiSCRIBE can be regulated both spatially and temporally, coupled to cellular regulatory circuits, and performed in a completely autonomous fashion, all of which provide greater ease and flexibility in adaptive writing and evolution experiments.

In summary, our work sheds light onto various factors that modulate the efficiency of retroelement-mediated recombineering and circumvents some of the limitations imposed by the exiting oligo-mediated recombineering and DNA writing systems in bacteria, offers a framework for the dynamic engineering of bacterial genomes with high efficiency and precision, and demonstrates and foreshadows multiple useful applications that are enabled by efficient and dynamic *in vivo* DNA writing. We envision that HiSCRIBE, along with the analogous DNA writing technologies demonstrated in eukaryotes (Anzalone et al., 2019; Sharon et al., 2018), will have broad utility in biotechnological and biomedical applications, including single-cell memory and computing, *in situ* engineering of genomes within communities, spatiotemporal molecular recording and connectome mapping, continuous *in vivo* evolution of single-gene (e.g., protein function) or multi-gene (e.g., metabolic network) traits, evolutionary engineering, and gene resurrection studies.

## Acknowledgments

We thank the MIT BioMicroCenter for technical support with next-generation sequencing. This work was supported by the National Institutes of Health (P50 GM098792), the Office of Naval Research (N00014-13-1-0424), the National Science Foundation (MCB-1350625), the Defense Advanced Research Projects Agency, the MIT Center for Microbiome Informatics and Therapeutics, and NSF Expeditions in Computing Program Award 1522074. F.F. would like to thank the Schmidt Science Fellows Program, in partnership with the Rhodes Trust for their support.

## Author contributions

F.F. and T.K.L. conceived the study. F.F. designed and performed the experiments. F.F. and N.G. designed experiments and analyzed next-generation sequencing data. R.J.C. contributed expertise with the phagemid experiments and edited the manuscript. F.F., N.G., and T.K.L. analyzed data and wrote the manuscript.

## Supplementary Materials

### 1. Supplementary Text

#### A Model for HiSCRIBE DNA Writing

Knocking out cellular exonucleases increases the background recombinant frequency in the uninduced HiSCRIBE system (Fig. 2A). This could be due to the leakiness of the promoter expressing the HiSCRIBE operon (P*_lacO_*) and/or an elevated recombination rate between the double-stranded DNA plasmid template and its target site in the Δ*recJ* Δ*xonA* background, as reported previously (Dutra et al., 2007). To investigate these two possibilities, we measured the recombinant frequency in the presence and absence of reverse transcriptase (RT) activity in different genetic backgrounds. An elevated recombinant frequency was observed even in the presence of inactive RT (Fig. S1A). However, in all of the tested conditions, cells expressing an active RT showed about two orders of magnitude greater recombinant frequencies compared to those expressing an inactive RT (Fig. 2A and Fig. S1A).

These results are consistent with a previously proposed model in which double-stranded plasmid templates can be recombined into a genomic target site via ssDNA intermediates through a RecA-independent process normally repressed by cellular exonucleases (Dutra et al., 2007). We speculate that even in the absence of an active retron system, recombinogenic oligonucleotides are produced *in vivo*, likely due to plasmid degradation by cellular nucleases. This intracellular ssDNA pool could then be processed and further degraded by cellular exonucleases, thus limiting the efficiency of recombination in the WT background. However, when cellular exonucleases (*recJ* and *xonA*) are knocked out, the intermediate degradation products of retron-encoded ssDNAs, as well as the template double-stranded DNA, could accumulate and contribute to the intracellular ssDNA pool, thereby increasing recombination efficiency (Fig. S1B). This model is further supported by previous observations in which the efficiency of oligo-mediated recombination directly correlated with the concentration of transformed oligos (Murphy and Marinus, 2010; Sawitzke et al., 2011). The addition of non-specific carrier ssDNAs can also compensate for low concentrations of specific ssDNA, potentially by transiently saturating cellular nucleases (Sawitzke et al., 2011). In this working model (Fig. S1B), Beta recombinase protects the intracellular oligonucleotide pool from cellular exonucleases and facilitates recombination between the ssDNAs and their corresponding genomic target loci.

In our experiments, ssDNAs were specifically designed to have at least three mismatches to the target in order to efficiently suppress the MMR system (Sawitzke et al., 2011) and achieve high-efficiency writing. Inefficient recognition of mismatched lesions, which is likely to occur in the absence of ssDNA expression in an exonuclease knockout background, could also contribute to the increased background observed in the HiSCRIBE system.

Knocking out *xseA*, which encodes one of the two subunits of ExoVII, slightly reduced the recombination efficiency (Fig. 2A). ExoVII is an ssDNA-specific exonuclease that converts large ssDNA substrates into smaller oligonucleotides (Chase and Richardson, 1974) and has been shown to be responsible for the removal of phosphorothioated nucleotides from the flanking ends of recombineering oligos (Mosberg et al., 2012), as well as the removal of the *msr* moiety from the msDNA of RNA-less retrons (Jung et al., 2015). Based on these observations, we speculate that ExoVII, among other cellular factors, may be involved in generating recombinogenic ssDNA intermediates. It is also possible that RecBCD-mediated processing of double-stranded breaks could provide another source for the intracellular recombinogenic ssDNA pool (Dillingham and Kowalczykowski, 2008).

Lastly, the optimal length of the flanking ssDNA homology arms that result in maximal HiSCRIBE editing efficiency was found to be around 35 bps (Fig. S1C). Increasing the size of the homology arm to 80 bp reduced the recombination efficiency, which we speculate could be due to secondary structures that prevent efficient recombination and/or inefficient ssDNA production by the retron system. These results are consistent with previous reports for recombineering with synthetic oligos (Sawitzke et al., 2011), and further confirm the involvement of a RecA-independent, Beta-mediated process in DNA writing by HiSCRIBE.

#### Measuring HiSCRIBE DNA Writing Efficiency with a Screenable Phenotype and High-throughput Sequencing

To systematically assess HiSCRIBE writing efficiency in an entire population, we used a screening assay with colorimetric readout. We introduced two stop codons into the *galK* ORF of the MG1655 Δ*recJ* Δ*xonA* reporter strain, hereafter referred to as *exo*^-^ *galK*_OFF_ strain. These reporter cells were transformed with HiSCRIBE(*galK*)_ON_ (HiSCRIBE plasmid encoding ssDNA identical to the WT *galK*). These cells were recovered for one hour in LB (37 C, 300 RPM) and plated on MacConkey + galactose (gal) + antibiotic plates in order to select for transformants. The conversion of the *galK*_OFF_ allele to *galK*_ON_ (i.e., the WT allele) was monitored by scoring the color of transformant colonies. As shown in Fig. S3, all the *galK*_OFF_ (white) cells transformed with the HiSCRIBE(*galK*)_ON_ plasmid formed galactose-fermenting *galK*_ON_ (pink) colonies on the indicator plates. No pink colonies were detected when cells were transformed with a non-specific HiSCRIBE [HiSCRIBE(NS)] plasmid. These results demonstrate that in the entire population of cells that received the HiSCRIBE(*galK*)_ON_ plasmid, *galK*_OFF_ alleles were converted to *galK*_ON_ over the course of colony growth, resulting in a phenotypic change in colony color.

Since Beta-mediated recombineering is a replication-dependent process (Huen et al., 2006; Murphy and Marinus, 2010), the conversion of *galK*_OFF_ to *galK*_ON_ occurs over the course of growth of the colonies, and a single pink colony observed on a transformation plate may contain a heterogeneous population of both edited and non-edited alleles. We measured the frequency of these alleles within single colonies by PCR amplification of the *galK* locus followed by Sanger sequencing as well as high-throughput sequencing. To avoid any difference in fitness between the two alleles in the presence of galactose, after we transformed the HiSCRIBE(*galK*)_ON_ plasmid into *exo*^-^ *galK*_OFF_ reporter cells, we selected transformants on LB plates, instead of MacConkey + gal plates. Sanger sequencing of PCR amplicons of the *galK* locus obtained from these transformants showed a mixture of peaks in the target site, suggesting that each colony on these plates may have contained a mixture of edited and non-edited alleles (Fig. S3). To give the replication-dependent HiSCRIBE writing system additional time to work, we re-streaked the colonies on fresh plates. Sanger sequencing of *galK* locus amplicons obtained from these colonies indicated the full conversion of the *galK*_OFF_ allele to *galK*_ON_, to the extent that the *galK*_OFF_ allele was below the limit of detection (Fig. S3). These amplicons were further quantified by high-throughput sequencing (Fig. S3). These results further validated Sanger sequencing results and indicated that HiSCRIBE system can be used to edit a desired genomic locus up to homogeneity (∼100% efficiency) in an entire population, and without the requirement for any double-strand DNA breaks and *cis*-encoded elements on the target.

#### Delivering HiSCRIBE via Different Strategies for Editing Bacteria within Bacterial Communities and Editing Non-traditional Hosts

To facilitate the delivery of HiSCRIBE for DNA writing in non-modified hosts, we placed the HiSCRIBE and CRISPRi systems into a single synthetic operon as shown in Fig. 3C and S4A, cloned it into a high-copy-number plasmid, and assessed its performance in the MG1655 *galK*_OFF_ reporter strain, which harbors two stop codons within the *galK* locus. Cells were chemically transformed with either HiSCRIBE(*galK*)_ON_ or HiSCRIBE(NS), which expressed a *galK_ON_* ssDNA or a non-specific ssDNA, respectively. The cells were recovered in LB for an hour, then plated on MacConkey + gal + antibiotic plates to select for HiSCRIBE plasmid delivery and screen for *galK_OFF_* to *galK_ON_* editing. More than 99% of cells transformed with the HiSCRIBE(*galK*)_ON_ plasmid formed pink colonies on these plates, indicating successful writing in the *galK* locus in all cells that received this plasmid (Fig. S4A). No pink colonies were detected in the samples transformed with the HiSCRIBE(NS) plasmid. The frequency of editing within individual colonies was assessed by PCR amplification of *galK* locus followed by high-throughput sequencing at 24 hours after transformation, as well as after a re-streaking step as described before (Fig. S4A).

Similar to transduction, conjugation is a common strategy for horizontal gene transfer in natural bacterial communities. In addition to using transduction for delivering HiSCRIBE plasmids (Fig. 3C), we tested whether conjugation can be used to deliver and edit cells within a complex bacterial community. We encoded the origin of transfer from RP4 (oriT) into the HiSCRIBE(*galK*)_ON_ plasmid and then introduced this plasmid into MFDpirPRO cells (that harbor RP4 conjugation machinery) to produce a donor strain. We showed that these cells could conjugate the HiSCRIBE(*galK*)_ON_ plasmid into recipient cells (MG1655 Str^R^ *galK*_OFF_). More than 99% of transconjugants formed pink colonies on MacConkey + gal + antibiotic plates (Fig. S4B), while no pink colonies were obtained in recipients that had been conjugated with the non-specific HiSCRIBE(NS) plasmid. We then conjugated the HiSCRIBE(*galK*)_ON_ plasmid into a stool-derived bacterial community containing MG1655 Str^R^ *galK*_OFF_, analogously to the transduction experiments (Fig. 3C). More than 99% of transconjugants that received the HiSCRIBE(*galK*)_ON_ plasmid formed pink colonies on the screening plates and no pink colonies were detected in cells conjugated with the non-specific HiSCRIBE(NS) plasmid (Fig. S4B). However, the efficiency of delivery via conjugation was significantly lower than phagemid transduction (Fig. S4C). We anticipate that more specific transduction delivery mechanisms are better suited for editing specific species within a community, while the more general (albeit less efficient) conjugation delivery mechanism is better suited for situations where editing a larger subpopulation of bacteria in the community are desired.

### 2. Materials and Methods

#### 2.1 Strains and Plasmids

Conventional cloning methods, Gibson assembly (Gibson, 2011) and Golden Gate assembly (Engler and Marillonnet, 2014) were used to construct the plasmids. Lists of strains and plasmids used in this study are provided in Tables S2 and S3, respectively. The sequences for the synthetic parts and primers are provided in Tables S4 and S5, respectively. Constructs will be available on Addgene.

#### 2.2 Cells and Antibiotics

Chemically competent *E. coli* DH5α F’ *lacI^q^* (NEB) was used for cloning. Unless otherwise noted, antibiotics and small molecule inducers were used at the following concentrations: Carbenicillin (Carb, 50 μg/mL), Kanamycin (Kan, 20 μg/mL), Chloramphenicol (Cam, 30 μg/mL), Streptomycin (Str, 50 μg/mL), Spectinomycin (Spe, 100 μg/mL), anhydrotetracycline (aTc, 200 ng/mL) and Isopropyl β-D-1-thiogalactopyranoside (IPTG, 1mM).

#### 2.3 Experimental Procedure

##### Induction of Cells and Plating Assays

The *kanR* reversion assay was performed as described previously (Farzadfard and Lu, 2014). Briefly, for each experiment, single colony transformants were separately inoculated into LB broth + appropriate antibiotics and grown overnight (37°C, 300 RPM) to obtain seed cultures. Unless otherwise noted, inductions were performed by diluting the seed cultures (1:1000) in LB + antibiotics ± inducers followed by 24 h (corresponding to log_2_(1000) ∼10 generations of growth) incubation (37°C, 700 RPM) in 96-well plates. Cultures were then serially diluted and spotted on selective media to determine the number of recombinant and viable cells in each culture. The number of viable cells was determined by plating serial dilutions of the cultures on LB plates with antibiotics corresponding to the marker present on the HiSCRIBE plasmid (Carb or Cam). LB + Kan plates were used to determine the number of recombinants. For each sample, the recombinant frequency was reported as the mean of the ratio of recombinants to viable cells for three independent replicates.

In the *galK* conversion assays, HiSCRIBE plasmids were delivered to reporter cells (with either chemical transformation, transduction or conjugation) and cells were recovered in LB for one hour without selection and plated on LB + appropriate antibiotics for HiSCRIBE plasmid selection. Allele frequencies were measured by MiSeq sequencing of colonies obtained on these plates after 24 h (corresponding to log_2_(10^9^) ∼30 generations of growth(Milo et al., 2010)). Additionally, for *galK*_OFF_ to *galK*_ON_ reversion experiments, cells were plated on MacConkey agar base (without carbon source) + galactose (1%) + appropriate antibiotics (for HiSCRIBE plasmid selection). The ratio of pink colonies (*galK*_ON_) to transformants (pink + white colonies) was used as a measure of recombinant frequency. For each sample, the recombinant frequency was reported as the mean of the ratio of recombinants to viable transformants for three independent replicates.

In the CRISPR-Cas9 counter-selection experiment (Fig. 3B), a gRNA against the *galK*_OFF_ locus (gRNA(*galK*_OFF_)) was placed under the control of an aTc-inducible promoter and cloned into the HiSCRIBE(*galK*)_ON_ plasmid. This plasmid was transformed into a *galK*_OFF_ reporter strain harboring an aTc-inducible Cas9 (or dCas9 as a negative control) plasmid. Single transformant colonies were diluted to ∼10^6^ CFU/mL in LB + Carb + Cam in the presence or absence of aTc and grown for 12 hours. These cultures were diluted and regrown for two additional cycles at the presence or absence of the inducer. The allele frequencies were determined by PCR amplification of the *galK* locus followed by high-throughput sequencing.

##### Phagemid Packaging

HiSCRIBE plasmids were packaged into M13 phagemid particles as described previously (Chasteen et al., 2006). Briefly, HiSCRIBE plasmids with the M13 origin of replication were transformed into an M13 packaging strain (DH5αPRO F^+^ harboring the M13cp helper plasmid) and the obtained single-colony transformants were grown overnight in 2 mL LB + antibiotics. The cultures were then diluted (1:100) in 50 mL fresh media and grown to saturation with selection. Phagemid particles were purified from the culture supernatants by PEG/NaCl precipitation (Yamamoto et al., 1970), passed through a 0.2-μm filter and stored in SM buffer (50 mM Tris-HCl [pH 7.5]), 100 mM NaCl, 10 mM MgSO_4_) at 4°C for later use.

##### Delivery by Transduction and Conjugation

For transduction experiments, overnight cultures of the reporter strain harboring an F-plasmid were diluted (1:1000) in fresh media and transduced by adding purified phagemid particles encoding HiSCRIBE at a Multiplicity of Infection (MOI) of 50, unless otherwise noted. After an hour incubation (37°C, 700 RPM), serial dilutions of the cultures were spotted on MacConkey + gal + antibiotics plates and recombinant frequency was calculated as described above (*galK* reversion assay).

For conjugation delivery, the MFDpirPRO strain was first produced by transforming the PRO plasmid (pZS4Int-*lacI*/*tetR*, Expressys) into the diaminopimelic acid (DAP)-auxotrophic MFDpir strain (Ferrieres et al., 2010) that encodes RP4 conjugation machinery. HiSCRIBE plasmids harboring RP4 origin of transfer were transformed into the MFDpirPRO strain to produce donor strains. Donor and recipient strains were grown overnight in LB with appropriate selection. Media for the donor strains was supplemented with 0.3 mM DAP throughout the experiment. In experiments shown in Figs. 4C and S4B, after conjugation, donor cells were selectively removed from conjugation mixtures by growing the cells in the absence of DAP.

Overnight cultures of donor and recipient strains were diluted (1:100) in fresh media and grown to an OD_600_ ∼1. Cells were pelleted and resuspended in LB, and mating pairs were mixed at a donor to recipient ratio of 100:1 and spotted onto nitrocellulose filters placed on LB agar supplemented with 0.3 mM DAP. The plates were incubated at 37°C for 6 h to allow conjugation. Conjugation mixtures were collected by vigorous vortexing the filters in 1 mL PBS, then serially diluted and spotted on MacConkey + gal + antibiotics plates as described in the *galK* reversion assay. The ratio of pink colonies per transconjugants was used as a measure of recombinant frequency.

For experiments showing genome editing in bacterial communities (Fig. 3C and S4B), an overnight culture of an undefined bacterial community was obtained by inoculating mouse stool in LB. This bacterial community was mixed (100:1) with a spontaneous Str^R^ resistant mutant of the MG1655 *galK*_OFF_ reporter strain to build a synthetic bacterial community that served as the recipient cell population in these experiments. For transduction experiments, the F plasmid was introduced to the reporter strain via conjugation using DH5α F^+^ (NEB) as the donor. The transduction and conjugation protocols were performed as described above, using the synthetic community as the recipient population.

##### Bacterial Connectome Mapping

To demonstrate that spatial information can be recorded into DNA memory, we mapped the pairwise connectome network of mating pairs in conjugating bacterial populations (Fig. 4C). The HiSCRIBE(Reg1)_r-rand_ library (overexpresses an ssDNA library with 6 randomized nucleotides targeting Register 1 in the *galK* locus, pZA11 backbone) was transformed into MG1655 Δ*recJ* Δ*xonA galK*_OFF_ to make a barcoded recipient population. A mobilizable HiSCRIBE(Reg2)_d-rand_ library (overexpressing an ssDNA library with 6 randomized nucleotides targeting Register 2 in the *galK* locus, pZE32 backbone) was transformed into MFDpirPRO cells to serve as the donor population. The donor and recipient populations were mixed at a 10:1 ratio (three parallel experiments) and conjugated as described above (37°C for 6 h). Conjugation mixtures were collected by vigorously vortexing nitrocellulose filters in 3 mL LB (without DAP) and recovered for 1 hour after which antibiotics (Carb + Cam) were added to select for transconjugants harboring HiSCRIBE(Reg1)_r-rand_ and HiSCRIBE(Reg2)_d-rand_ plasmids. Samples were grown at 37°C overnight in the absence of DAP to selectively remove donor cells and allow HiSCRIBE writing and propagation of the edited alleles. Genomic DNA was prepared from the overnight cultures and the contents of memory registers were analyzed by high-throughput sequencing as described below.

For the bacterial organization mapping experiment (Fig. 4B), barcoded clonal donor and recipient populations harboring HiSCRIBE(Reg1)_r-barcode_ and HiSCRIBE(Reg2)_d-barcode_ were spotted as indicated patterns and conjugated as described above (37°C for 6 h). After conjugation, allele-specific PCR was used (see below) to amplify the edited registers directly from conjugation mixtures (without any outgrow).

##### High-throughput Sequencing

Allele frequencies of the HiSCRIBE target sites were measured by sequencing amplicons obtained from corresponding genomic sites using Illumina MiSeq. Target loci were amplified using 1 μL of liquid culture (or colony resuspension) as a template. Barcodes and Illumina adapters were then added in an additional round of PCR. Samples were gel-purified, multiplexed, and sequenced by Illumina MiSeq. The obtained reads were demultiplexed based on the attached barcodes and mapped to the reference sequence.

For *galK* conversion experiments, any reads that lacked the expected “ATGCCXXXXXXATCGAT” motif, where “XXXXXX” corresponds to the 6-bp variable site in the *galK* alleles (TTGCTG for *galK*_WT_, CTATTA for *galK*_SYN_, CTCTTG for *galK*_ON_, and TAATGA for *galK*_OFF_), or that contained ambiguous nucleotides within this region were discarded. For *galK*_WT_ to *galK*_SYN_ experiment, editing efficiency was reported as the ratio of *galK*_SYN_ reads to the total number of *galK*_SYN_ + *galK*_WT_ reads. For *galK* reversion experiments, editing efficiency was calculated as the ratio of *galK*_ON_ reads to the total number of *galK*_ON_ + *galK*_OFF_ reads. The enrichment of recombinant alleles in the WT *E. coli* MG1655 background (Fig. S4A) was investigated similarly. Single colonies of transformants were picked 24 h (or 48 h) after transformation, resuspended in water, and used as templates for PCR. The samples were processed as described above.

A similar strategy was used to analyze the dynamics of the P*_lac_* locus in the experiment shown in Fig. 5. The P*_lac_* locus was amplified using 1 μL of liquid culture obtained from samples at different time points throughout the experiment. Barcodes and Illumina adapters were added in an additional round of PCR. Samples were gel-purified, multiplexed, and sequenced by paired-end Illumina MiSeq for higher accuracy. Any reads that lacked the expected “YYYYYYCTTTATGCTTCCGGCTCGZZZZZZ” motif, where “YYYYYY” and “ZZZZZZ” correspond to positions of the -35 and -10 boxes of the P*_lac_* promoter, respectively, or that contained ambiguous nucleotides within this region were discarded. The variant frequencies were calculated as the ratio of the number of reads for a given variant to the total number of reads for that sample.

For the bacterial spatial organization recording and connectome mapping experiments (shown in Fig. 4B and Fig. 4C, respectively), barcoded donor and recipient populations were conjugated as described above. For the former experiment, conjugation mixtures were resuspended in LB and the memory registers in the *galK* locus were amplified by allele-specific PCR to deplete unedited registers (which mainly originate from cells that did not undergo successful conjugation, which form the majority of conjugation mixtures). As shown in Fig. S5A, we designed primers that specifically bind to the writing control nucleotide of edited alleles but form a mismatch (at the 3’-end position) with the unedited registers. We then used these primers and HiDi DNA polymerase (a selective variant of DNA polymerase that can only amplify templates that are perfectly matched at the 3’-end with a given primer, myPLOS Biotec, DE) to specifically amplify edited registers from 1 μL of conjugation mixtures while depleting the unedited registers. Illumina barcodes and adapters were then added to the samples by a second round of PCR. Samples were gel-purified, multiplexed, and sequenced by Illumina MiSeq. Samples were then computationally demultiplexed, and any reads that contained non-edited registers, which lacked any of the two expected motifs flanking the two memory registers (ATGCCTMMMMMMTCGATT and AGTGCGNNNNNNGTGCGC, where “MMMMMM” and “NNNNNN” correspond to positions of the memory Registers 1 and 2, respectively), or that contained ambiguous nucleotides within this region were discarded. The frequencies of variants that were observed simultaneously in a single read in the two registers were then calculated and presented as weighted connectivity matrices (Figs. 4B and S5B).

For the latter experiment, an alternative depletion strategy was used. Specifically, genomic DNA was purified from overnight cultures of the conjugation mixtures using the ZR Fungal/Bacterial DNA MiniPrep kit (Zymo Research). A DNA fragment including Registers 1 and 2 in the *galK* locus was PCR amplified from purified genomic DNA and gel purified. The samples were depleted of non-edited (i.e., WT) sequences by enzymatic digestion with ClaI and AgeI, since these sites are present in non-edited Register 1 and 2, but are removed after HiSCRIBE recording. Samples were subsequently run on TBE gels (6%) and uncut fragments (edited in both Registers) (Fig. S6A) were extracted for purification. Mixed sequence populations were detected in the two memory registers by Sanger sequencing, indicating successful writing in both registers (Fig. S6B). Illumina barcodes and adapters were added to the purified sample by a second round of PCR followed by enzymatic digestion as described above to remove residual non-edited registers. Samples were gel-purified, multiplexed, and sequenced by Illumina MiSeq (300 bps, single-end). Any reads that contained non-edited registers, that lacked any of the two expected motifs flanking the two memory registers (ATGCCTMMMMMMTCGATT and AGTGCGNNNNNNGTGCGC, where “MMMMMM” and “NNNNNN” correspond to positions of the memory Registers 1 and 2, respectively), or that contained ambiguous nucleotides within this region were discarded. The connectivity matrices were deduced by linking variants that were observed simultaneously in a single read in the two registers and presented as heatmaps. To capture as many interactions as possible, we used an inclusive approach and did not filter out infrequent reads, which could potentially result in false positives due to the relatively high error rate of MiSeq. As an additional control, and in order to estimate the false-positive discovery rates due to sequencing errors or spontaneous mutations, we calculated a connectivity matrix for two randomly chosen (non-targeted) 6-bp regions within the *galK* amplicon. Only a limited number of connections were detected (Fig. S6C and Supplementary File S1). Further inspection of these mutated non-targeted regions revealed that they were mostly comprised of single base pair differences with the wild-type sequences, suggesting that these arose from sequencing errors, which are reportedly ∼10^-3^-10^-2^ mutations per nucleotide (Ross et al., 2013). False positives could be further reduced by using error-reducing library preparations, computational correction methods, and/or more accurate sequencing platforms (Lou et al., 2013; Ross et al., 2013; Schmitt et al., 2012).

##### Continuous Evolution of the P*_lac_* Promoter

The efficient genome editing achieved by HiSCRIBE can be coupled with continuous selection or screening to enable the continuous evolution of desired target loci. In order to demonstrate this adaptive writing strategy, we chose to evolve P*_lac_* in *E. coli* (Fig. 5). To achieve a wider dynamic range of fitness, we started with a weakened P*_lac_* promoter, created by mutating the -10 sequence of P*_lac_* promoter from “TATGTT” to “CCCCCC”. This mutation leads to poor growth of cells in M9 media when lactose is the sole carbon source. An overnight culture of the parental strain harboring the mutated P*_lac_* promoter (MG1655 Δ*recJ* Δ*xonA* F^+^ P*_lac_*_(“TATGTT”→“CCCCCC”)_) was diluted (1:100) into M9 + glu (0.2%) and divided into two groups, each with three parallel cultures. Samples in each group were treated with phagemid particles (MOI = 100), from either a HiSCRIBE(P*_lac_*) phagemid library or the non-specific [HiSCRIBE(NS)] control, and incubated in a microplate reader at 37°C with continuous shaking (250 RPM). The cultures were grown for 1 hour before antibiotic selection (Carb and Cam for phagemid delivery and F-plasmid maintenance, respectively). Cells were then grown for 23 additional hours, diluted (1:100) into M9 + lactose (0.2%) + phagemid + antibiotics, and grown for 48 hours at 37°C in a microplate reader as above. The dilution and regrowth (24 h) cycles were repeated five additional times to permit the selection and propagation of beneficial mutations. OD_600_ was monitored and samples were taken for Illumina sequencing throughout the experiment. Population growth rates based on OD_600_ were calculated using the GrowthRates tool (Hall et al., 2014).

To verify the activity of the identified variants in the P*_lac_* evolution experiments, we reconstructed these variants in the parental background using oligo-mediated recombineering (Chan et al., 2007). The reconstructed variants were grown overnight in LB, diluted (1:100) in fresh media supplemented with IPTG (1 mM), and grown for 8 hours (37°C, 700 RPM). The activities of reconstructed P*_lac_* promoter variants were measured by Miller assay using Fluorescein di-β-D-galactopyranoside (FDG) as the substrate. 50 μL of each culture was mixed with 50 μL of B-PER II reagent (Pierce Biotechnology) containing FDG (0.005 mg/mL final concentration). The fluorescent signal (absorption/emission: 485/515 nm) was monitored in a plate reader with continuous shaking for 2 hours at 37°C. β-galactosidase activity was calculated by normalizing the rate of FDG hydrolysis (obtained from fluorescence signal) to the initial OD_600_. For each sample, β-galactosidase activity was reported as the mean of three independent biological replicates.

##### HiSCRIBE Library Construction

Randomized HiSCRIBE phagemid and mobilizable libraries (for experiments shown in Figs. 4C and 5, respectively) were constructed by a modified Quik-Change (Agilent) protocol. Briefly, HiSCRIBE plasmids (with or without the RP4 origin of transfer) were PCR amplified using primers containing the randomized regions within the desired target site in the overhangs. The primers also contained compatible sites for the type IIS enzyme Esp3I. PCR products were used in a Golden Gate assembly (Engler and Marillonnet, 2014) using this cut site to circularize the vector amplicon. Circularized vector libraries were amplified by transformation into Electroten Blue electrocompetent cells (Agilent). Amplified libraries were then packaged into phagemid particles for transduction experiments (as described above) or transformed into donor and recipient strains and used in the mating pair connectome mapping experiment as described above.

---

**Figure S1.**
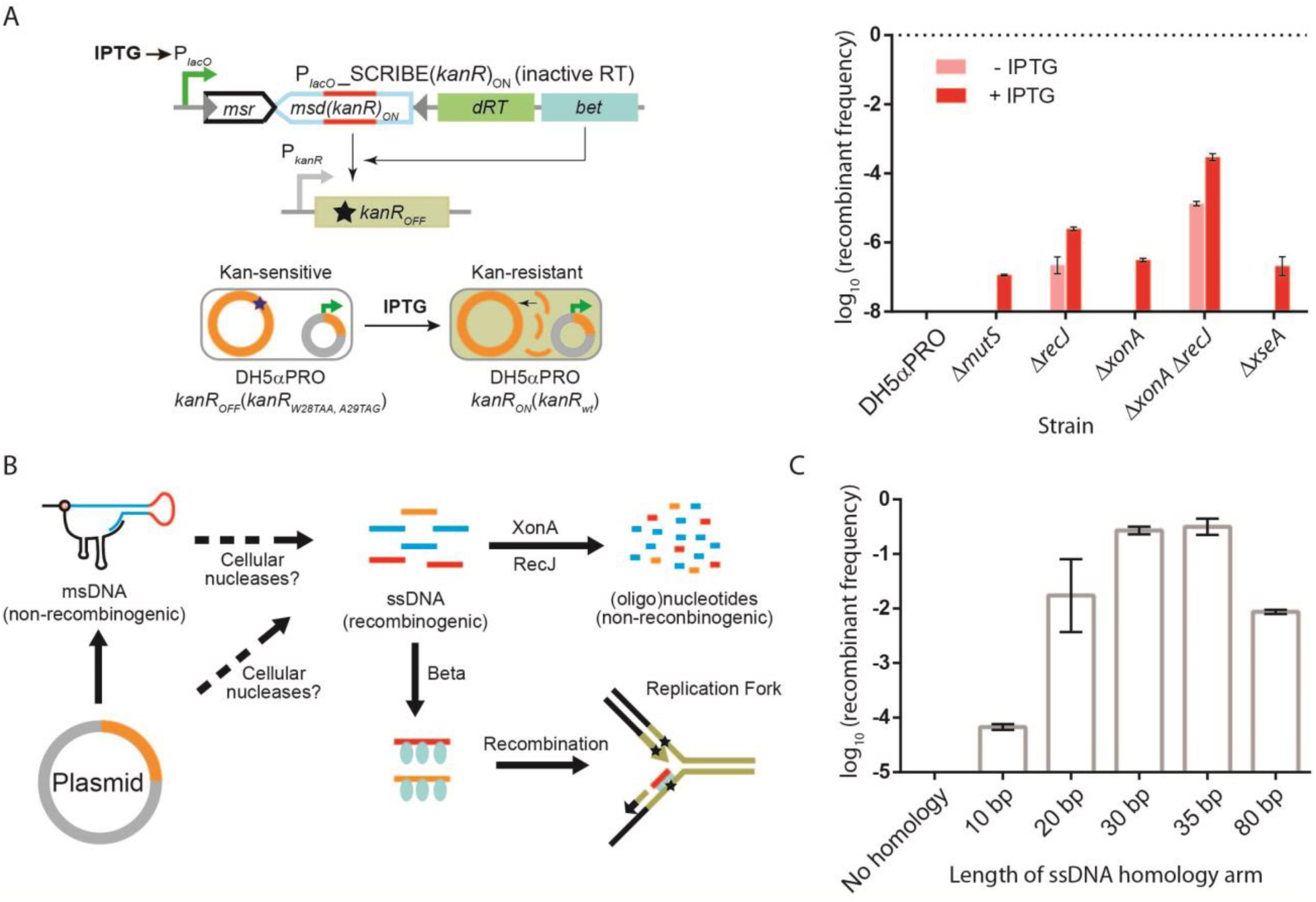
A model for HiSCRIBE-mediated recombineering. **(A)** Genome editing efficiencies of SCRIBE harboring a catalytically inactive reverse transcriptase (*dRT*, in which the conserved YADD motif in the active site of the RT is replaced with YAAA (Farzadfard and Lu, 2014)) was determined by the *kanR* reversion assay in different knockout backgrounds. Error bars indicate standard error of the mean for three biological replicates. **(B)** Proposed model for retron-mediated recombineering. Intracellular recombinogenic oligonucleotides are likely generated due to the degradation of the template plasmid as well as msDNA (retron product). ssDNA-specific cellular exonucleases (XonA and RecJ) can process these oligonucleotides into smaller, non-recombinogenic (oligo)nucleotides. Alternatively, Beta can bind to, protect, and recombine these oligonucleotides into their genomic target loci. **(C)** Effect of ssDNA homology length on HiSCRIBE DNA writing efficiency. Different HiSCRIBE(*kanR*)_ON_ plasmids expressing ssDNAs with various lengths of homology to the *kanR*_OFF_ target were tested by the *kanR* reversion assay in DH5αPRO Δ*recJ* Δ*xonA kanR*_OFF_ reporter strain. Maximal editing efficiency was observed with ssDNAs encoding 35 bp homology arms. Error bars indicate standard errors for three biological replicates.

**Figure S2.**
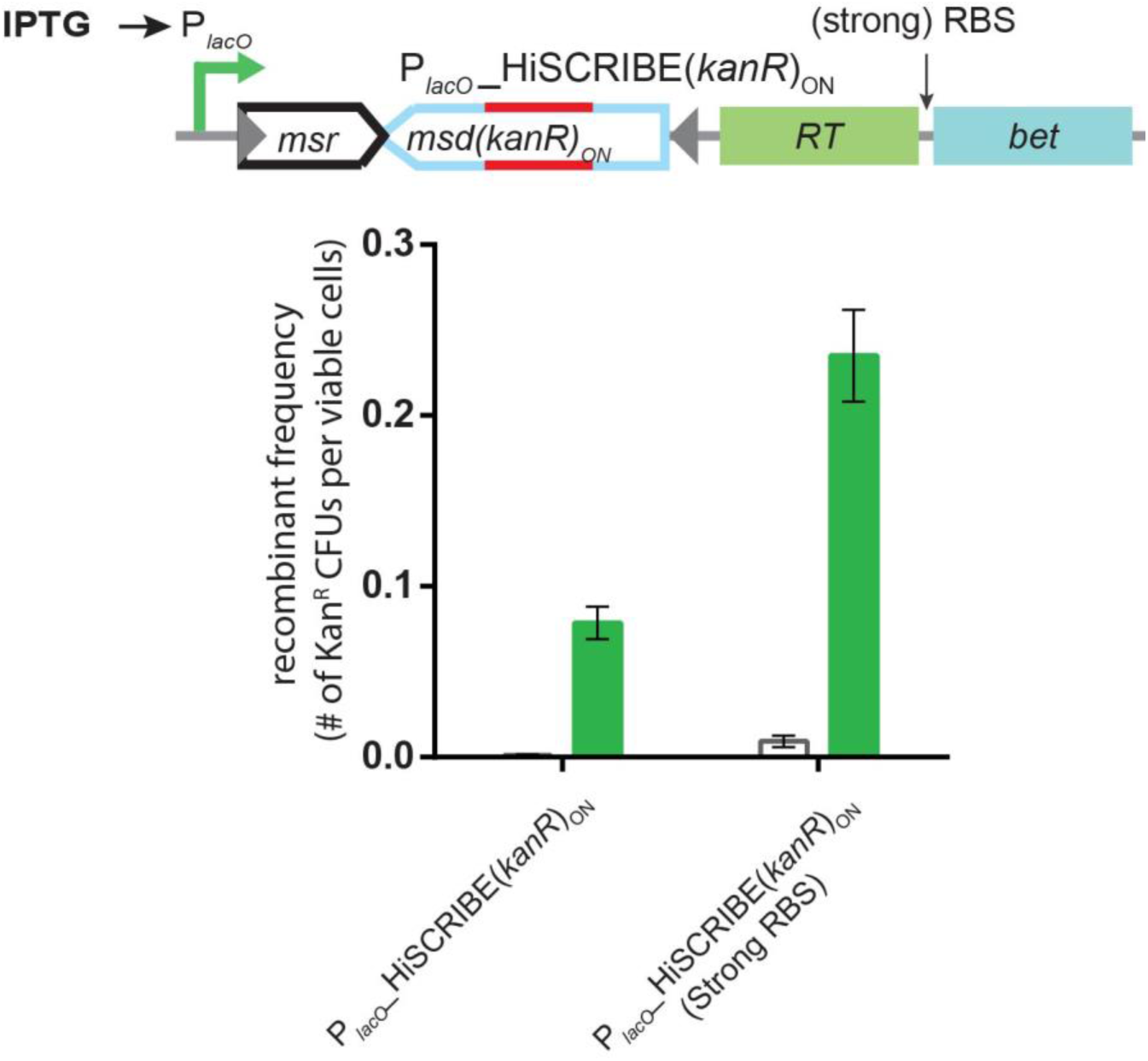
Optimizing HiSCRIBE efficiency by tuning the expression level of Beta. DH5αPRO Δ*recJ* Δ*xonA kanR_OFF_* reporter cells were transformed with IPTG-inducible HiSCRIBE(*KanR*)_ON_ constructs, harboring either natural *bet* RBS or a strong synthetic RBS (Zelcbuch et al., 2013), and the recombinant frequency was measured using the *kanR* reversion assay. Error bars indicate standard errors for three biological replicates.

**Figure S3.**
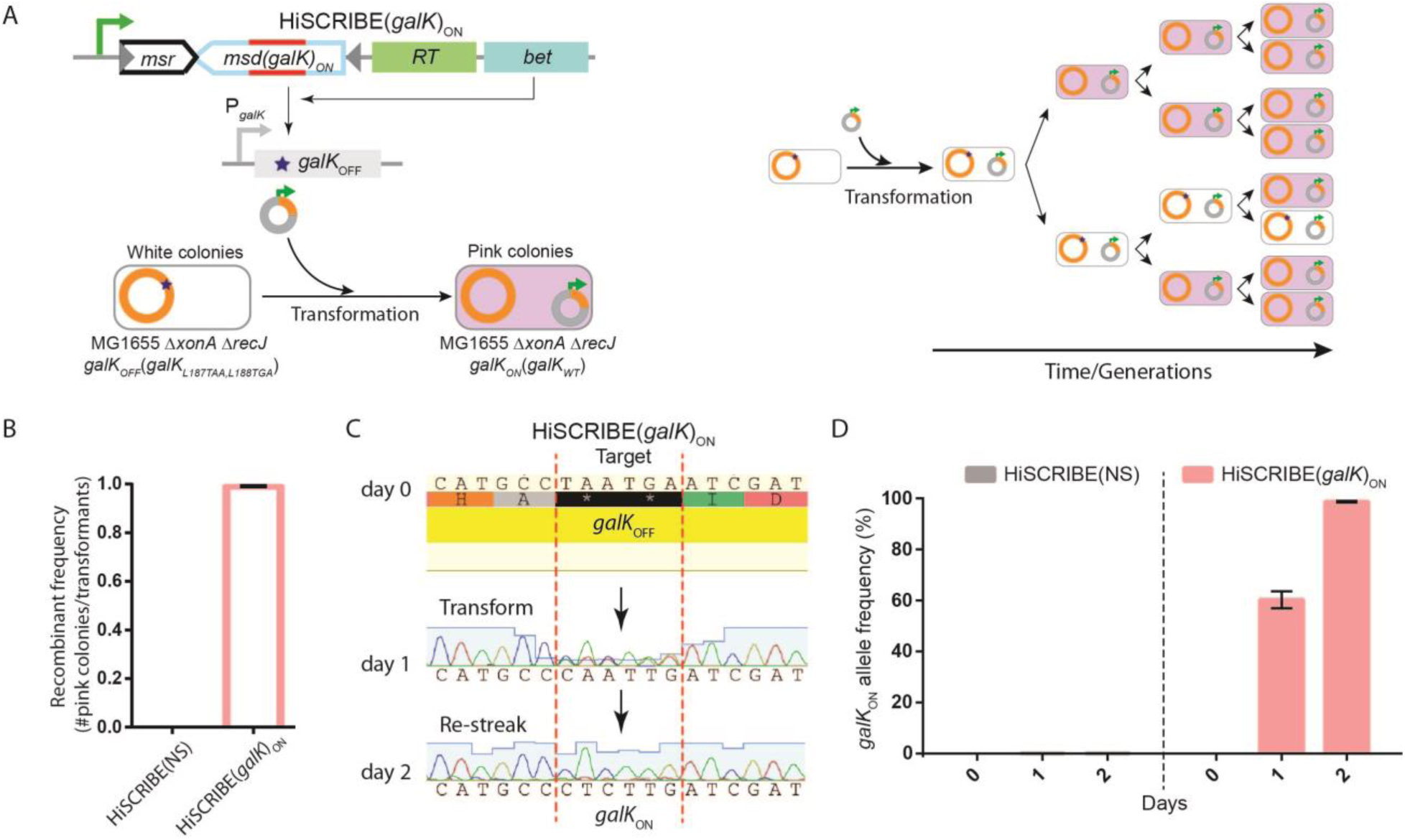
Assessing population-wide HiSCRIBE writing efficiency using plating assay and sequencing. **(A)** The genetic circuit used to assess writing efficiency (left panel) as well as the schematic representation of the enrichment of mutant alleles within a single transformant colony (right panel). **(B)** MG1655 *exo^-^ galK*_OFF_ reporter cells were transformed with the HiSCRIBE(*galK*)_ON_ plasmid and population-wide recombinant frequency was measured by the *galK* reversion assay. The frequencies of *galK*_ON_ and *galK*_OFF_ alleles in individual transformant colonies obtained on LB plates were assessed one and two days after transformation using **(C)** Sanger sequencing as well as **(D)** high-throughput Illumina sequencing.

**Figure S4.**
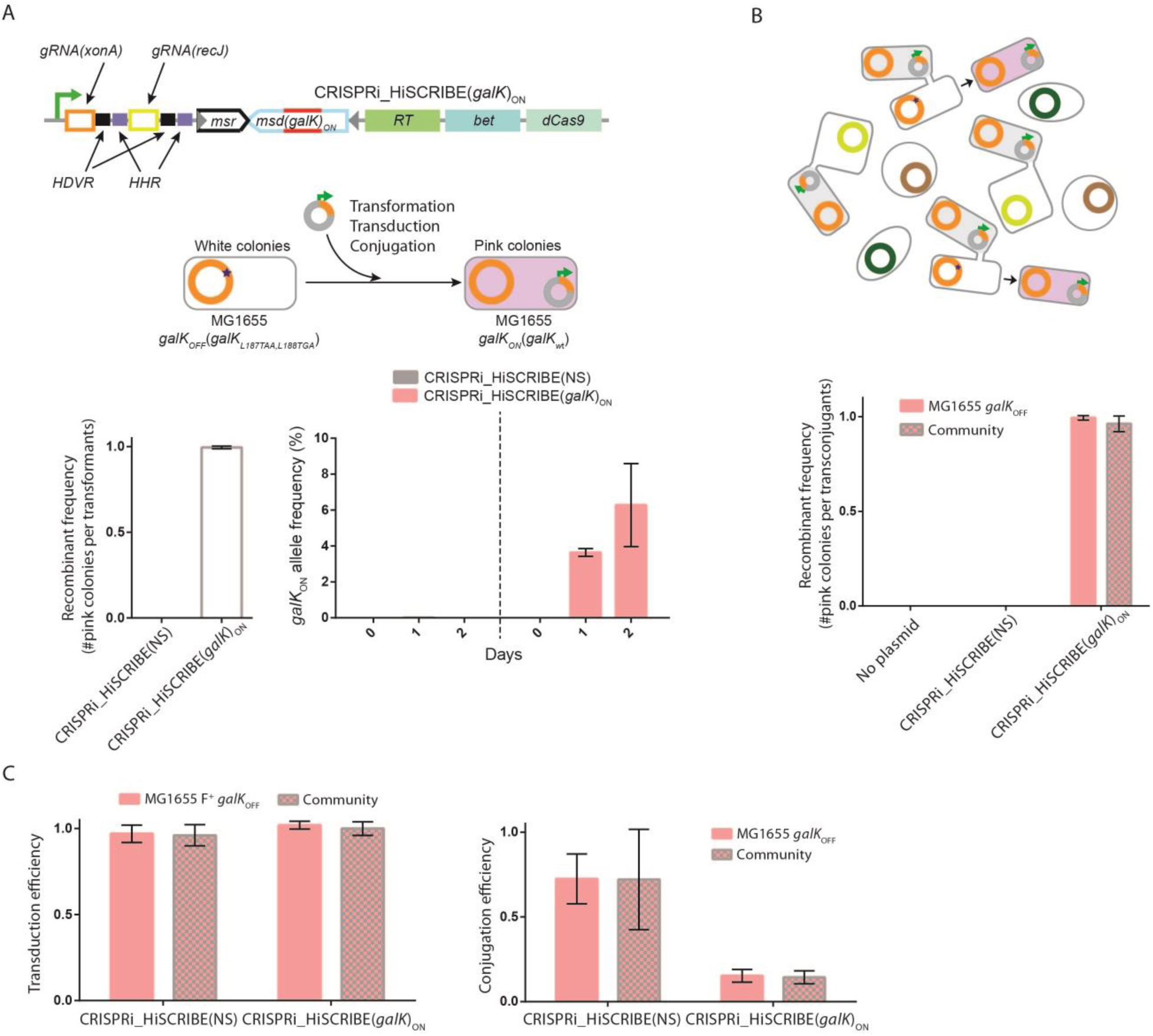
Efficient editing of bacterial genomes in clonal populations as well as within bacterial communities. **(A)** HiSCRIBE(*galK*)_ON_ was cloned into a ColE1 plasmid encoding both the M13 origin and RP4 origin of transfer and delivered into the MG1655 *galK*_OFF_ reporter strain via chemical transformation, transduction, and conjugation. Recombinant frequencies in cells that received HiSCRIBE(*galK*)_ON_ or HiSCRIBE(NS) by chemical transformation were assessed using the *galK* reversion assay. Allele frequencies of individual transformant colonies obtained on LB with appropriate selection were measured by Illumina sequencing 24 hours after transformation, as well as after 24 hours of additional growth. **(B)** Using a conjugative HiSCRIBE plasmid (harboring RP4 origin of transfer) to edit the MG1655 galK*_OFF_* Str^R^ reporter strain in the clonal population as well as within a synthetic bacterial community. **(C)** The delivery efficiency of HiSCRIBE plasmid by transduction and conjugation (for the experiments shown in Fig. 3C and S4B, respectively). To assess the transduction efficiency of HiSCRIBE phagemids, transduction mixtures were serially diluted and plated on LB + Str and LB + Str + Carb plates, to measure the number of viable target cells and transductants, respectively. The ratio between the transductants and viable target cells was reported as transduction efficiency. To measure the conjugation efficiency of delivering the HiSCRIBE plasmids, conjugation mixtures were serially diluted and plated on LB + Str and LB + Str + Carb plates, to measure the number of viable target cells and transconjugants, respectively. The ratio between the transconjugants and recipient cells was reported as conjugation efficiency.

**Figure S5.**
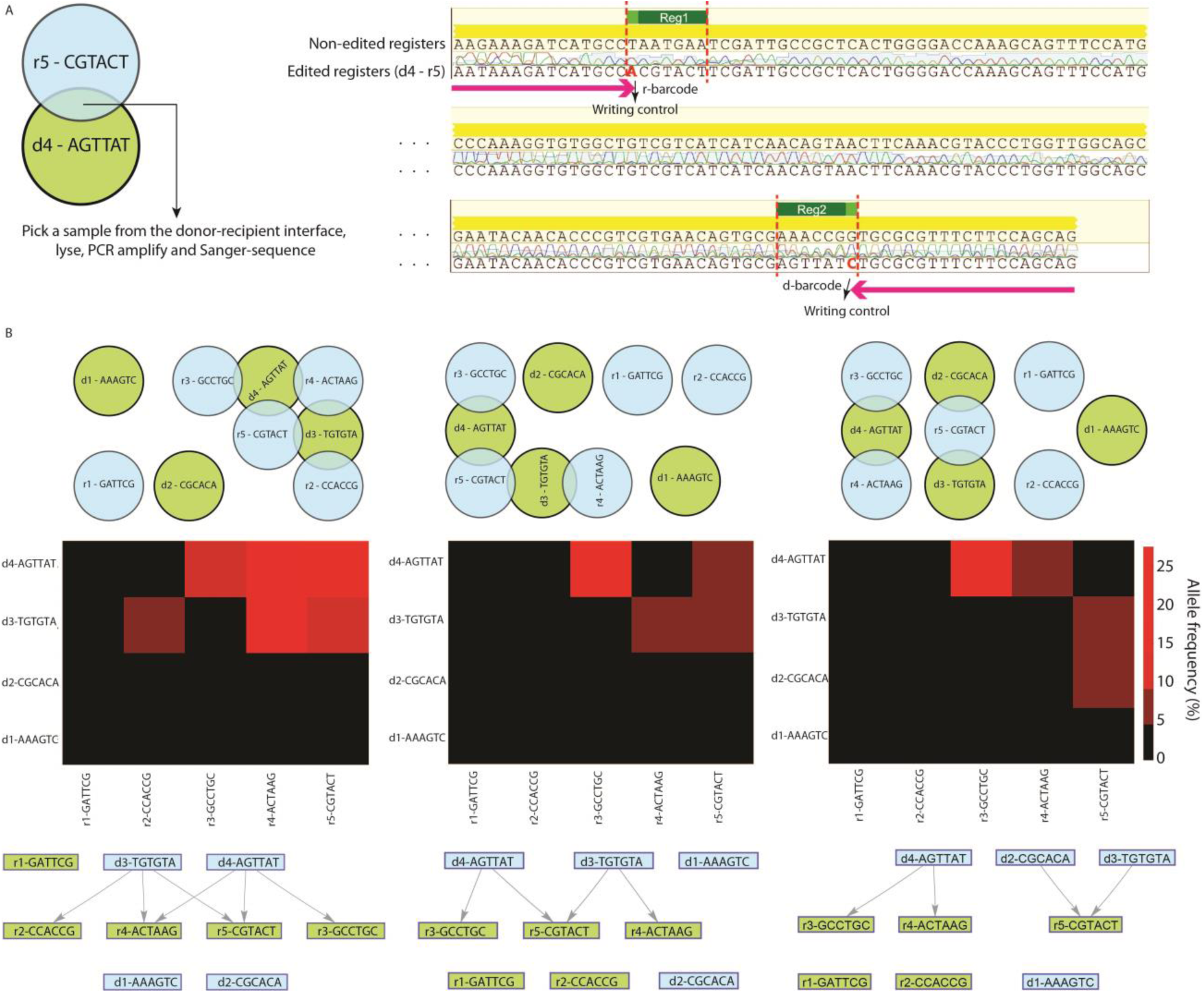
Recording spatial patterns of mating pairs in bacterial cell populations into DNA. **(A)** Conjugation donor and recipient populations harboring HiSCRIBE-encoded “*d-barcode*” and “*r-barcode*” were spotted on nitrocellulose filters placed on the agar surface as indicated by the green and blue circles, respectively. These plasmids were designed to introduce unique 6 bp barcodes, as well as additional mismatches (which serve as “writing control nucleotides” to discriminate between edited and unedited memory registers when selectively PCR amplifying the edited registers) into two adjacent memory register on the *galK* locus, once inside the recipient cells. Samples taken from the intersection of the donor and recipient populations were lysed and used as templates in allele-specific PCR. Allele-specific PCR using primers that bind to the “writing control nucleotides” (but not to the non-edited registers) was used to selectively amplify the edited registers and deplete non-edited registers. The identities of the two barcodes corresponding to the interacting donor and recipient populations were then retrieved by Sanger sequencing. **(B)** Additional examples of cellular patterns that were recorded by the barcode joining approach described in Fig. 4A and 4B, and their corresponding weighted connectivity matrices and interaction networks that were faithfully retrieved using high-throughput sequencing.

**Figure S6.**
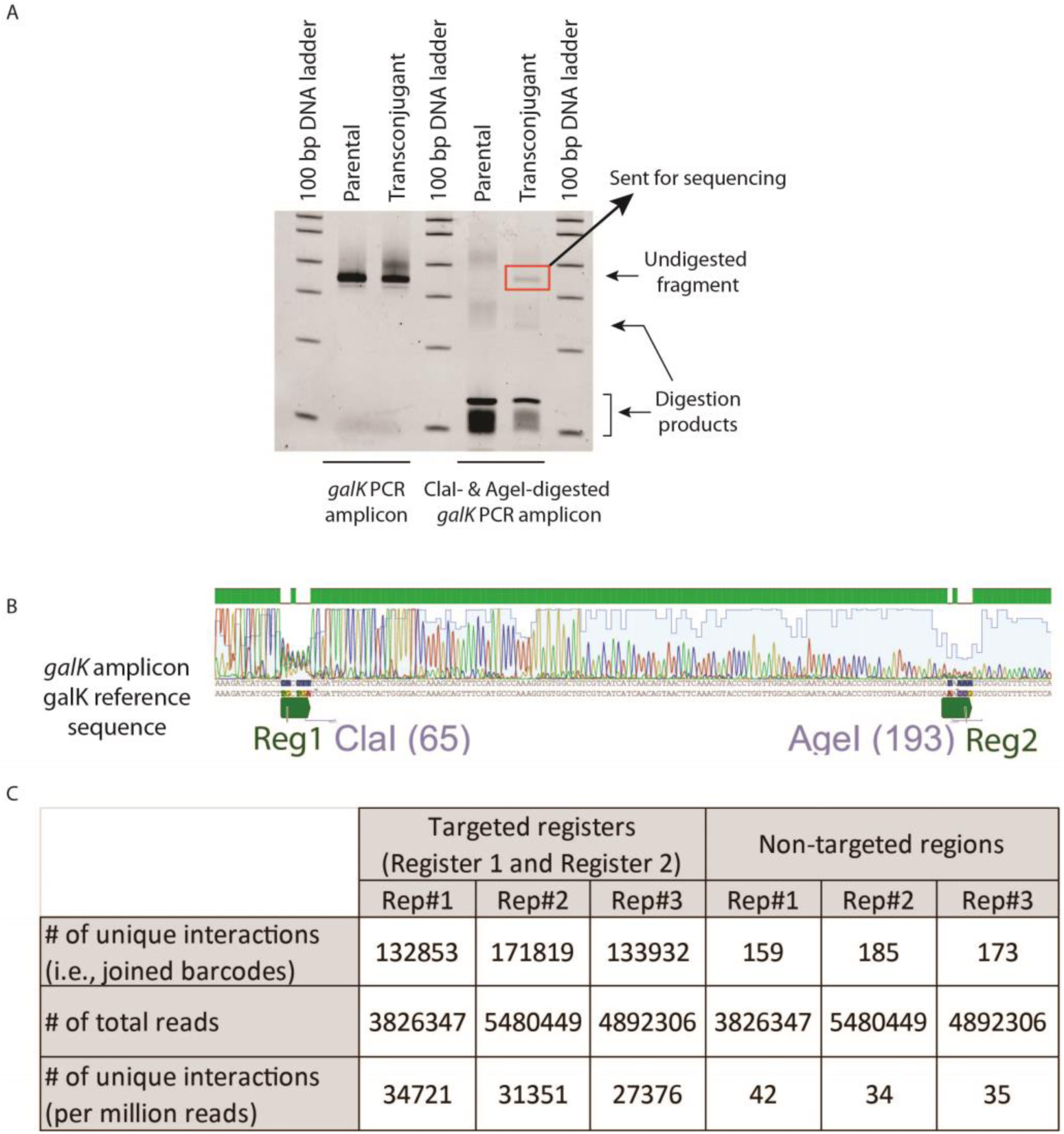
Strategy used to deplete unedited memory registers from dual-register amplicons and the frequency of cell-cell interactions recovered by high-throughput sequencing in the connectome mapping experiment. **(A)** Using restriction digestion as an alternative strategy to remove unedited registers from the PCR amplified amplicons instead of allele-specific PCR. Genomic DNA samples were purified from the parental recipient cells (MG1655 Δ*recJ* Δ*xonA galK*_OFF_), as well as cultures obtained after conjugation (transconjugants) in the experiment described in Fig. 4C. The *galK* locus was PCR amplified from the purified genomic DNA samples and run on a 6% TBE gel before and after digestion with ClaI and AgeI enzymes (which cut unedited Register 1 and Register 2, respectively) and stained by SYBR gold. The *galK* amplicon obtained from the parental sample was completely digested after enzymatic digestion. In contrast, the *galK* amplicon obtained from the transconjugant sample was not completely digested by ClaI and AgeI. The undigested band, corresponding to edited registers, comprised ∼3.9% of the signal in this lane (measured by densitometry). **(B)** This band was subsequently excised, purified and Sanger-sequenced. Drops in the quality of sequencing in Register 1 and 2 indicate the presence of mixed DNA populations containing variations in these two regions in these samples. Subsequently, Illumina adaptors and barcodes were added to this undigested amplicon using an additional round of PCR and the obtained amplicon was sequenced by Illumina MiSeq (see Methods). **(C)** Number of unique variants (interactions) per million reads obtained from sequencing the two target registers in the genomes of recipient cells after conjugation with donor cells, as well as two randomly selected non-targeted regions within the *galK* amplicon (used as a negative control and to assess the rate of false-positives), for the experiment shown in Fig. 4C.

**Figure S7.**
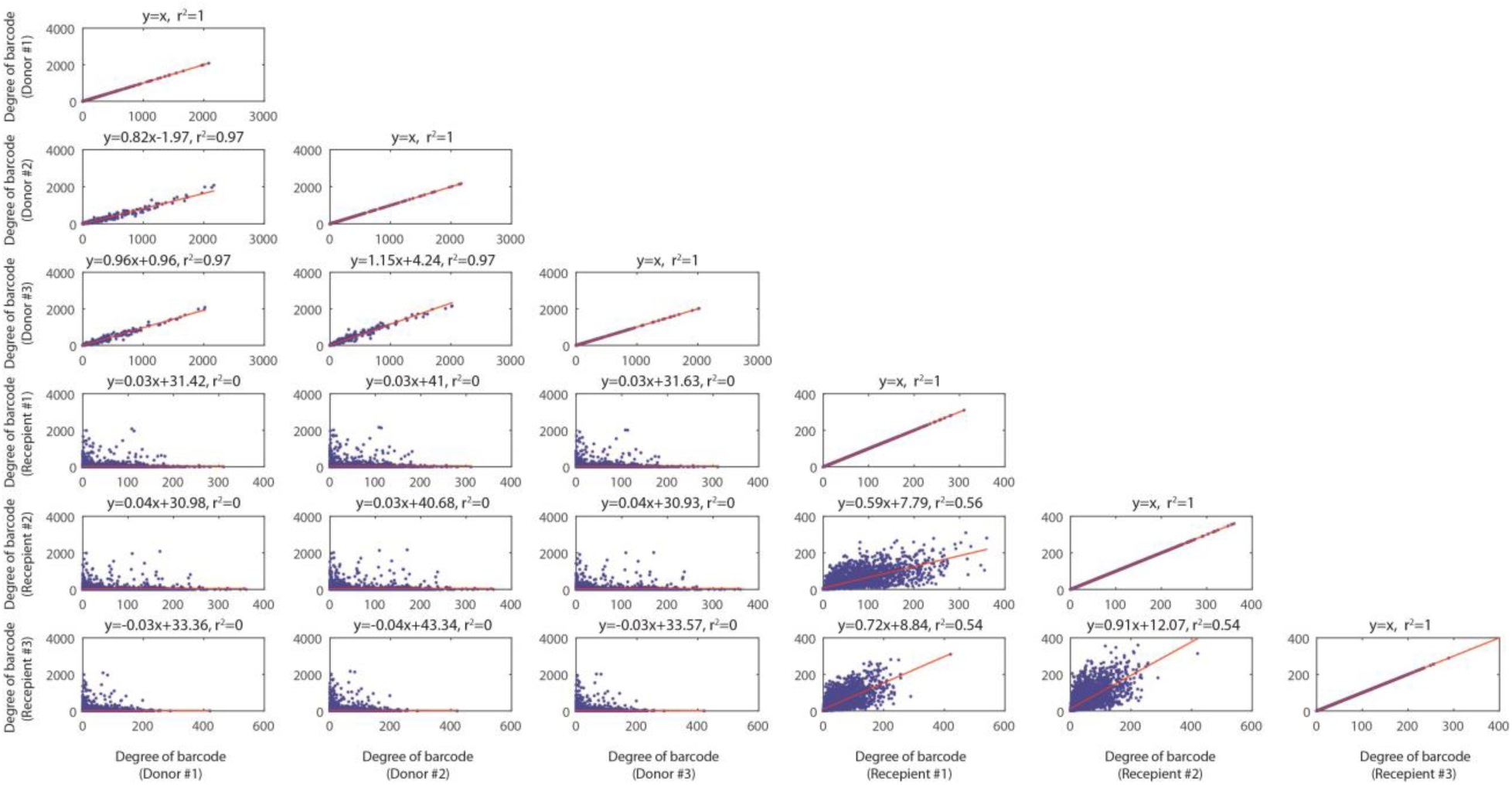
Correlation between degree of nodes for donor and recipients for three parallel conjugation mixtures. Correlations between degrees of donor barcodes and degrees of recipient barcodes for three parallel conjugation experiments. The degree of donor barcodes is defined as the number of unique interactions that each donor barcode makes with recipient barcodes, which is equal to the sum of elements of the column corresponding to that barcode in the presented connectivity matrix. The degree of recipient barcodes is defined as the number of unique interaction that each recipient barcode makes with donor barcodes, which is equal to the sum of elements of the row corresponding to that barcode in the presented connectivity matrix. The strong correlation between the degree of donor barcodes in the parallel conjugation experiments suggests that the transfer of barcodes from donors is not dependent on the identity of their conjugation partners (i.e., recipients). On the other hand, the relatively weak correlation between the degree of recipient barcodes suggests that other factors, such as the identities of the partners recipient cells partners (i.e., donor cells), might affect the frequency of successful conjugation at the tested donor:recipient ratio.

**Figure S8.**
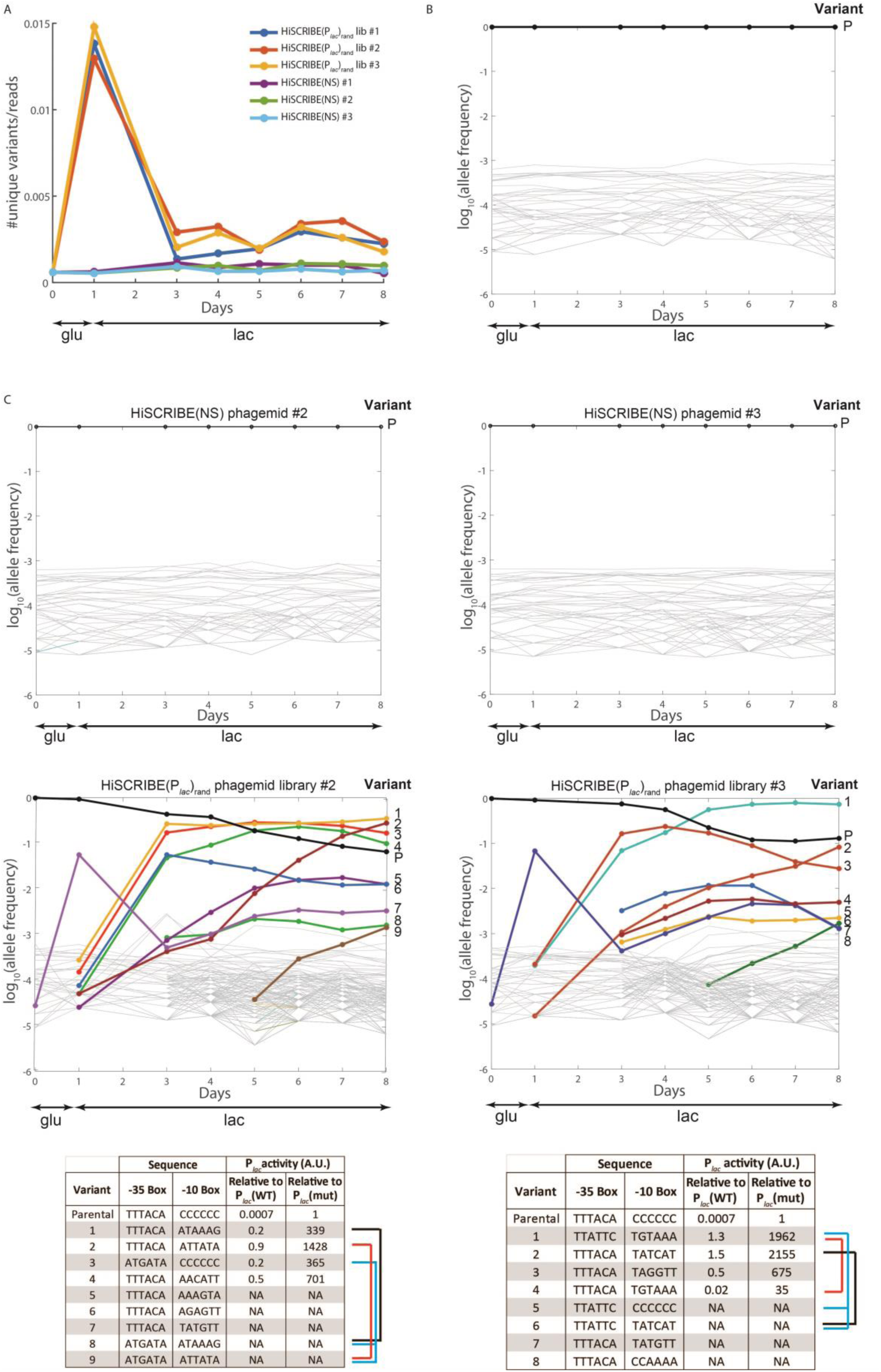
Dynamics of P*_lac_* alleles in the P*_lac_* evolution experiment. **(A)** The diversity of P*_lac_* alleles observed in the evolution experiment shown in Fig. 5 as well as two additional parallel cultures, reported as the number of unique variants per sequencing read. The diversity of the P*_lac_* locus in cultures exposed to the HiSCRIBE(P*_lac_*)_rand_ phagemid library was significantly higher than those exposed to HiSCRIBE(NS) phagemids. **(B)** Dynamics of P*_lac_* alleles for cultures that were exposed to HiSCRIBE(NS) phagemids in the experiment shown in Fig. 5. **(C)** Changes in P*_lac_* alleles frequencies over the course of the experiment shown as time series for cells exposed to the HiSCRIBE(NS) (top) or the HiSCRIBE(P*_lac_*)_rand_ library phagemid particles (middle) for two additional parallel cultures of the experiment shown in Fig. 5. The identities of the most frequent alleles at the end of the experiment, as well as fold-change in the β-galactosidase activity of the corresponding allele compared to the WT and parental alleles, are shown in the bottom tables. Alleles that are likely ancestors/descendants are linked by brackets.

**Table S1.** Side-by-side comparison of different features of currently available DNA writing systems in bacteria.

|  | HiSCRIBE<br>(this work) | Oligo-mediated<br>recombineering<br>(e.g. MAGE)<br>(Costantino and<br>Court, 2003;<br>Wang et al.,<br>2009) | Cas9-nuclease<br>assisted<br>genome<br>editing (Jiang<br>et al., 2013) | Base editing<br>(Farzadfard et<br>al., 2019;<br>Gaudelli et al.,<br>2017a; Komor<br>et al., 2016) | PRIME editing<br>(Anzalone et<br>al., 2019)<br>Only<br>demonstrated<br>in mammalian<br>cells | Cas1-Cas2<br>spacer<br>acquisition<br>(Shipman<br>et al., 2016) | Site-specific<br>recombinases<br>(Roquet et al.,<br>2016; Siuti et<br>al., 2013) |
| --- | --- | --- | --- | --- | --- | --- | --- |
| Requires<br>presence of *cis*-<br>elements on the<br>target | No | No | Yes (PAM) | Yes (PAM and<br>dC or dA<br>residues within<br>editing window) | Yes (PAM) | Yes<br>(CRISPR<br>array leader<br>sequence<br>and<br>repeats) | Yes |
| Requires<br>introduction of<br>dsDNA breaks | No | No | Yes | No | No | No | No |
| ~100% DNA<br>writing efficiency | Yes | No | Yes | Yes | No | No | Yes |
| Editing can be<br>linked to<br>biological events | Yes | No<br>(requires<br>exogenous<br>delivery of ssDNA<br>donors) | Yes (but<br>requires<br>delivery of DNA<br>donors) | Yes | Yes | Yes | Yes |
| Types of small<br>modifications<br>(insertions,<br>deletions, or<br>base-substitution<br>mutations) | Any | Any | Any | dC to dT<br>or dA to dG<br>or vice versa | Any | Small fixed-<br>size<br>insertions | Flipping or<br>excising DNA<br>located<br>between<br>recombinase<br>sites |

**Table S2.**
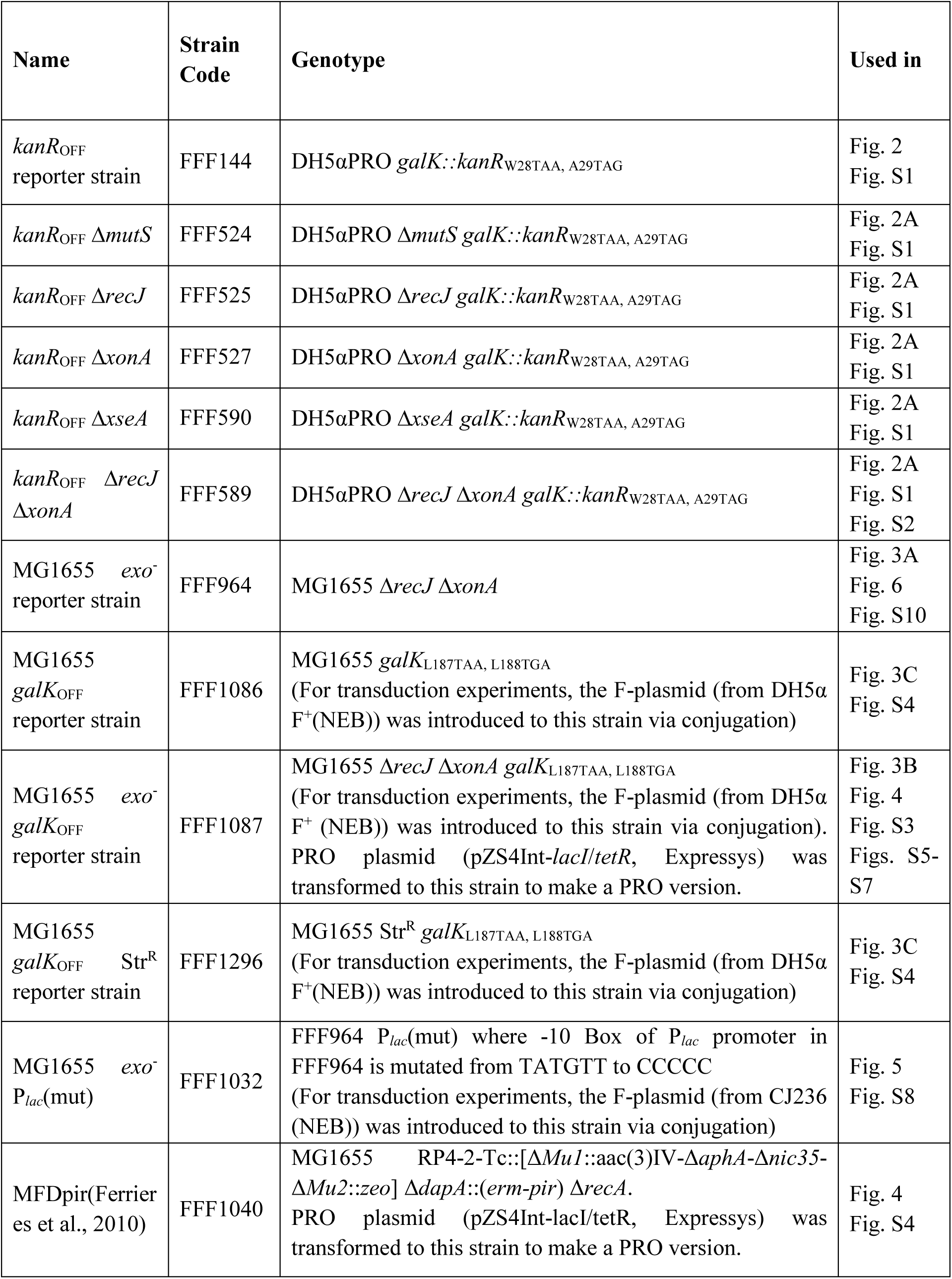
List of the reporter strains used in this study.

**Table S3.** List of the plasmids used in this study.

| Name | Plasmid Code | Maker | Used in | Ref |
| --- | --- | --- | --- | --- |
| PRO plasmid (pZS4Int-*lacI/tetR*) | pFF187 | Spe/Str | Fig. 3B<br>Fig. 4<br>Fig. S5-7 | Expressys (Lutz and Bujard, 1997) |
| pKD46 | pFF59 | Carb | Fig. 3C | (Datsenko and Wanner, 2000) |
| $P_{lacO\_msd}(kanR)_{ON}$ | pFF530 | Cam | Fig. S2 | (Farzadfard and Lu, 2014) |
| $P_{tetO\_bet}$ | pFF145 | Carb | Fig. S2 | (Farzadfard and Lu, 2014) |
| $P_{lacO\_SCRIBE}(kanR)_{ON}$ | pFF745 | Cam | Fig. 2A | (Farzadfard and Lu, 2014) |
| $P_{lacO\_SCRIBE}(kanR)_{ON\_dRT}$ | pFF755 | Cam | Fig. S1 | (Farzadfard and Lu, 2014) |
| $P_{lacO\_HiSCRIBE}(kanR)_{ON}$<br>(Strong RBS) | pFF804 | Cam | Fig. S2 | This work |
| $P_{lacO\_HiSCRIBE}(kanR)_{ON}$<br>(Strong RBS) | pFF944 | Carb | Fig. S1C | This work |
| $P_{tetO\_CRISPRi}$(no gRNA)<br>[or $P_{tetO\_dCas9}$] | pFF1156 | Cam | Fig. 2B<br>Fig. 3B | Addgene #44249<br>(Qi et al., 2013) |
| $P_{tetO\_CRISPRi}(recJ\_gRNA \& xonA\_gRNA)$ | pFF1165 | Cam | Fig. 2B | This work |
| $HiSCRIBE(galK)_{SYN}$<br>(Strong RBS) | pFF1493 | Carb | Fig. 3A | This work |
| $HiSCRIBE(galK)_{ON}$<br>(Strong RBS) | pFF1081 | Carb | Fig. S3 | This work |
| $HiSCRIBE(galK)_{ON\_P_{tetO\_gRNA}(galK_{OFF})}$ | pFF1220 | Carb | Fig. 3B | This work |
| $P_{tetO\_Cas9}$ | pFF1172 | Cam | Fig. 3B | This work |
| $CRISPRi\_HiSCRIBE(galK)_{ON}$ | pFF1298 | Carb | Fig. 3C<br>Fig. S4 | This work |

**Table S4.**
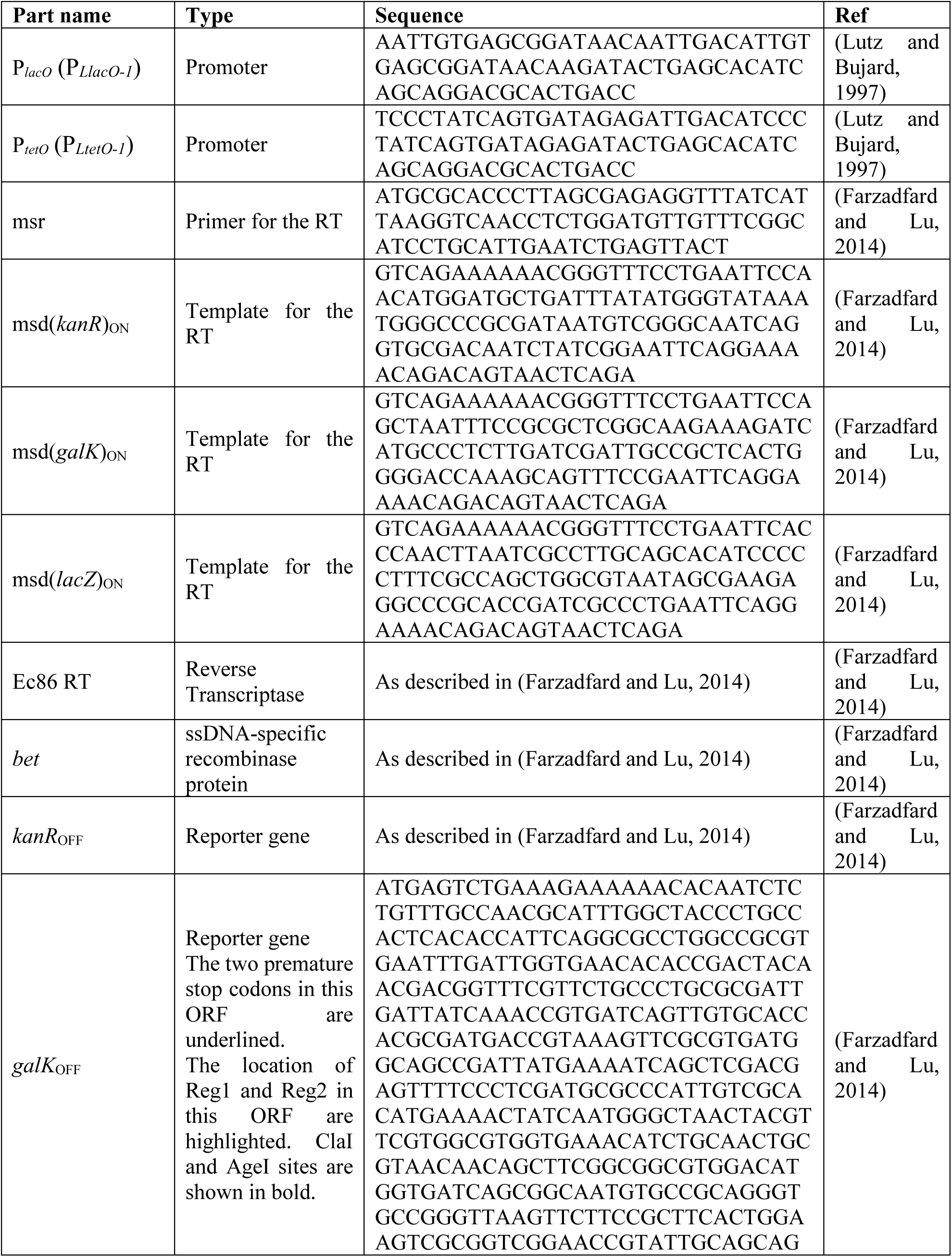
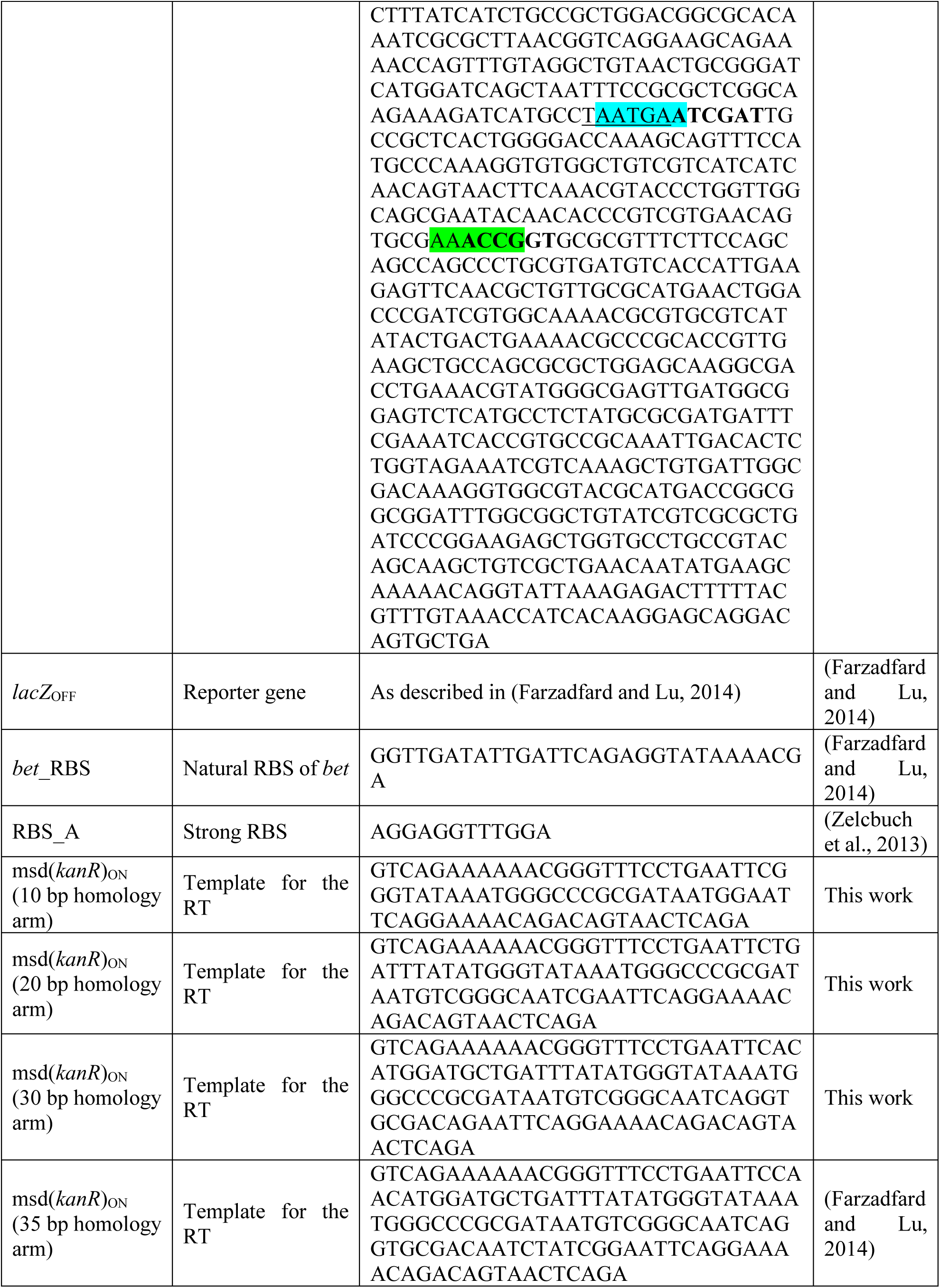
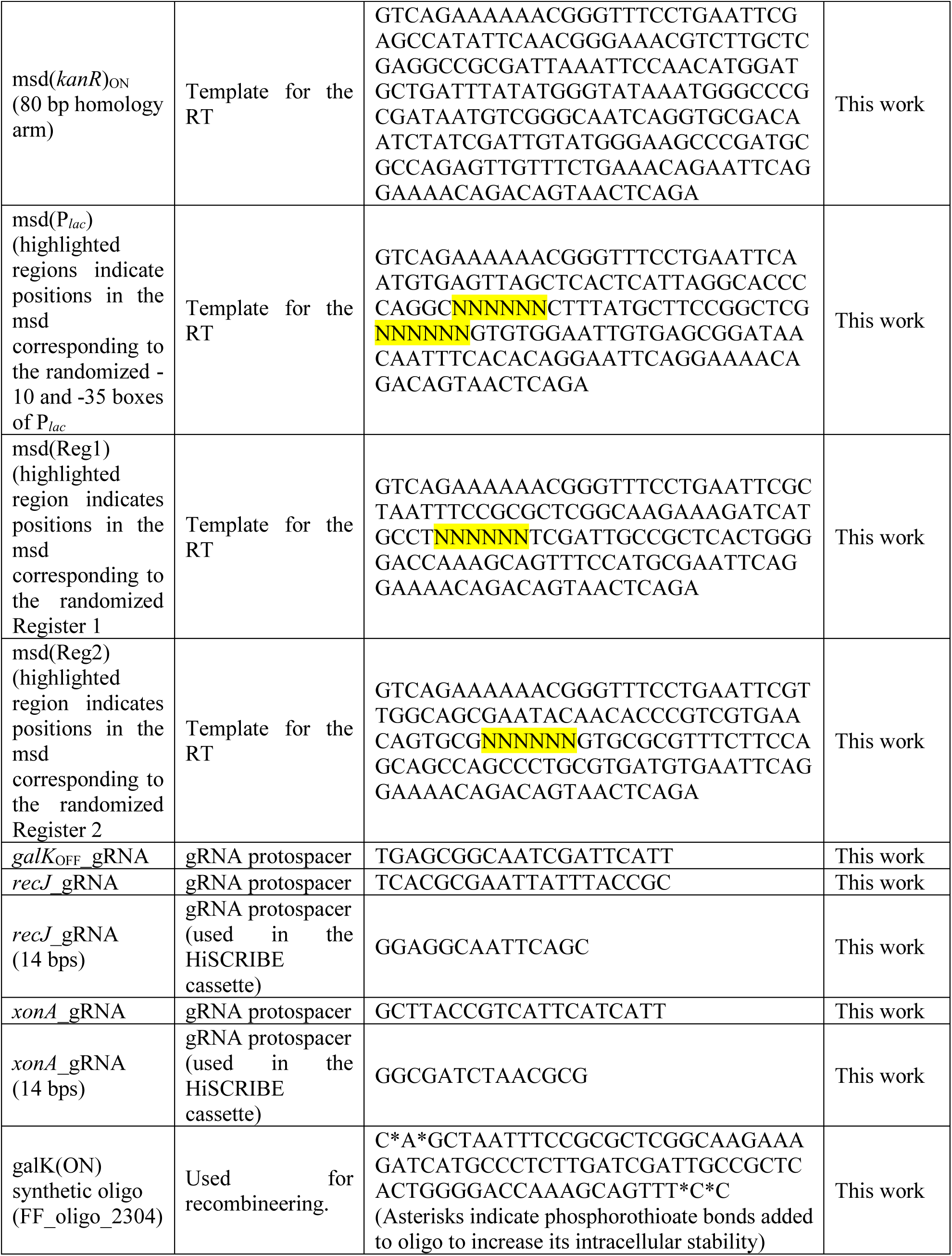
List of the synthetic parts and their corresponding sequences used in this study.

**Table S5.**
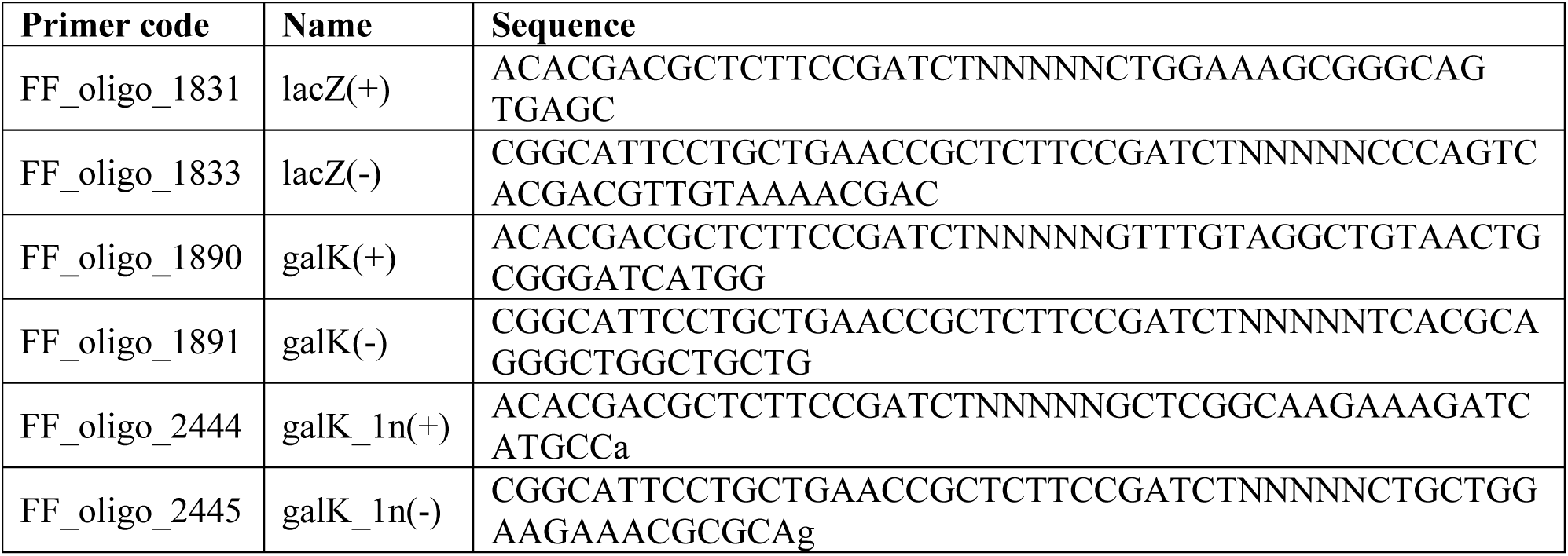
List of the sequencing primers used in this study.

